# Conserved secreted effectors determine endophytic growth and multi-host plant compatibility in a vascular wilt fungus

**DOI:** 10.1101/2021.08.29.457830

**Authors:** Amey Redkar, Mugdha Sabale, Christian Schudoma, Bernd Zechmann, Yogesh K. Gupta, Manuel S. López-Berges, Giovanni Venturini, Selena Gimenez-Ibanez, David Turrà, Roberto Solano, Antonio Di Pietro

**Affiliations:** Departamento de Genética, Universidad de Córdoba, 14071 Córdoba, Spain; Earlham Institute, Norwich Research Park, Colney Lane, Norwich, NR4 7UZ, United Kingdom; Baylor University, Center for Microscopy and Imaging, Waco, Texas 76798, USA; The Sainsbury Laboratory, Norwich Research Park, Colney Lane, Norwich, NR4 7UH, United Kingdom; Plant Molecular Genetics Department, Centro Nacional de Biotecnologıa-CSIC (CNB-CSIC), 28049 Madrid, Spain; Department of Agriculture and Center for Studies on Bioinspired Agro-enviromental Technology, Università di Napoli Federico II, 80055 Portici, Italy

**Author notes:** Correspondence: A. Redkar; A. Di Pietro. European Molecular Biology Laboratory (EMBL), Structural and Computational Biology Unit, Meyerhofstraße 1, 69117 Heidelberg, Germany. Isagro S.p.A. Centro Ricerche, Via Giacomo Fauser 28, 28100 Novara, Italy.

## Abstract

Fungal interactions with plant roots, either beneficial or detrimental, have a major impact on agriculture and ecosystems. The soil inhabiting ascomycete *Fusarium oxysporum* (Fo) constitutes a species complex of worldwide distribution causing vascular wilt in more than a hundred different crops. Individual isolates of the fungus exhibit host-specific pathogenicity, determined by proteinaceous effectors termed secreted in xylem (SIX). However, such isolates can also colonize roots of non-host plants asymptomatically as endophytes, or even protect them against pathogenic isolates. The molecular determinants of multi-host plant colonization are currently unknown. Here, we identified a set of fungal effectors termed ERCs (<u>E</u>arly <u>R</u>oot <u>C</u>ompatibility effectors), which are secreted during early biotrophic growth of Fo on both host and non-host plants. In contrast to the strain-specific SIX effectors, which are encoded on accessory genomic regions, ERCs are encoded on core regions and are found across the entire Fo species complex as well as in other phytopathogens, suggesting a conserved role in fungus-plant associations. Targeted deletion of ERC genes in a pathogenic Fo isolate resulted in reduced virulence on the host plant and rapid activation of plant immune responses, while in a non-pathogenic isolate it led to impaired root colonization and loss of biocontrol ability. Strikingly, some ERCs also contribute to Fo infection on the non-vascular land plant *Marchantia polymorpha*. Our results reveal an evolutionarily conserved mechanism for multi-host colonization by root infecting fungi.

## Introduction

Pathogenic microorganisms have evolved to infect both aerial as well as below-ground organs resulting in devastating losses of agricultural yield (1). Some pathogenic species target the host vasculature resulting in systemic infections, while others remain restricted to nonvascular tissues, causing localized disease symptoms. These interactions are the result of distinct adaptations allowing pathogens to accommodate themselves inside the plant host and dampen the immune response through secreted molecules termed effectors (2). Effectors function either in the intercellular space (apoplast) or inside the host cells to reprogram plant processes (3). Plants in turn, have evolved a multi-layered immune system to resist the microbial invaders and thwart pathogen success (4, 5).

Vascular wilt fungi constitute a particularly destructive group of soil-borne pathogens, which attack almost every crop except cereals and are extremely difficult to control (6). The *Fusarium oxysporum* (Fo) species complex provokes devastating losses in global agriculture(7, 8), exemplified by the highly aggressive clone named tropical race 4 (TR4) that threatens to wipe out the world’s industrial Cavendish banana production (9). Fo infection initiates in the soil, when the fungus senses chemical signals released by roots that trigger directed hyphal growth towards the plant (10). After entering the plant, Fo grows mainly between the cells of the root cortex. During this symptomless biotrophic stage, establishment of infection depends on conserved fungal compatibility mechanisms such as a mitogen-activated protein kinase cascade controlling invasive growth (11) or <u>R</u>apid <u>AL</u>kalinization <u>F</u>actor (RALF), a small secreted protein mimicking plant regulatory peptides that trigger host alkalinization (12). In a compatible pathogen-host interaction, Fo eventually enters and colonizes the xylem vessels, causing characteristic vascular wilt symptoms and plant death (7).

A long standing question concerns the molecular determinants of host range during the asymptomatic endophytic phase. Although the Fo species complex collectively infects more than a hundred different crops, individual isolates only cause wilt disease on a single or a few related plant species and have accordingly been classified into formae speciales (ff. spp.) (13, 14). Host-specific pathogenicity is conferred by accessory or lineage specific (LS) genomic regions, which encode unique combinations of virulence determinants and can be horizontally transferred between Fo isolates (14, 15). Many of these host-specific effector proteins were originally identified in the xylem sap of infected plants and are thus known as SIX (Secreted In Xylem) effectors (16-18). How SIX effectors determine host specificity is not fully understood, although some were shown to suppress host Pattern Triggered Immunity (PTI) and also function as avirulence genes in Effector Triggered Immunity (ETI) through recognition by host resistance (*R*) genes (7,19-21).

Besides causing wilt disease on their respective host species, Fo isolates can also colonize the roots of other plants where they survive asymptomatically as endophytes or even act as biocontrol agents to protect the plant against pathogenic Fo forms or other root pathogens (22-25). The genetic determinants underlying multi-host asymptomatic root compatibility in Fo remain unclear. In this study we investigated how Fo establishes multi-host root compatibility independent of its colonization lifestyle, be it pathogenic or endophytic. By analyzing the apoplastic fluid of infected roots, we identified a set of Early <u>R</u>oot <u>C</u>ompatibility (ERC) effectors, which are secreted during early biotrophic growth on both host and non-host plants and are found across the entire Fo species complex as well as in other fungal phytopathogens. Interestingly, ERCs also contribute to Fo infection on an evolutionarily distant non-vascular plant suggesting a broadly conserved role in fungus-root interaction.

## Results

### *F. oxysporum* colonizes roots of host as well as non-host plants

Fo isolates are known to colonize the roots of both host and non-host plants. At present the molecular basis of asymptomatic non-host invasion as opposed to vascular colonization and wilting remains largely unknown. To address this question, we first tested the ability of a set of Fo reference isolates sequenced by the Broad Institute Fusarium Comparative Genome Initiative (FCGI) to colonize a single plant species, tomato (table S1). The set includes several plant pathogenic ff. spp. infecting tomato, melon, pea, banana or crucifer species, as well as a non-pathogenic biocontrol isolate (24). All tested isolates were detected by real time quantitative PCR (qRT-PCR) in roots and root crowns of tomato plants at 35 days post inoculation (dpi), while only the tomato pathogenic isolate (f. sp. *lycopersici*) was found in the stems (fig. S1A). Confocal microscopy of tomato roots inoculated with fluorescent mClover3-labelled strains of the tomato pathogenic isolate Fol4287 (f. sp. *lycopersici*), the banana pathogenic isolate Foc54006 (f. sp. *cubense*) or the non-pathogenic biocontrol strain Fo47, confirmed the presence of fungal hyphae of all three strains which efficiently colonized the tomato root cortex (Fig. 1A). Moreover, transmission electron microscopy (TEM) of Fol4287 infected plants at 3 dpi revealed that the fungus localized predominantly to the apoplast of root cortical cells, although occasional penetration events of cortical and endodermal cells were observed, accompanied by a loss of plant plasma membrane integrity (Fig. 1B and fig. S1B). Again, qRT-PCR detected the presence of all three Fo isolates in tomato roots at 12 dpi, while only the tomato pathogenic isolate Fol4287 was detected in the stems (Fig. 1C). Collectively, these results confirm that Fo isolates have a general capacity to colonize roots of both host and non-host plants, which is in line with earlier reports (23). Our results also show that Fo colonization of non-host plants remains mostly restricted to roots and root crowns while the host-specific forms of the fungus are able to invade the stems to cause vascular wilt and plant death.

**Fig. 1.**
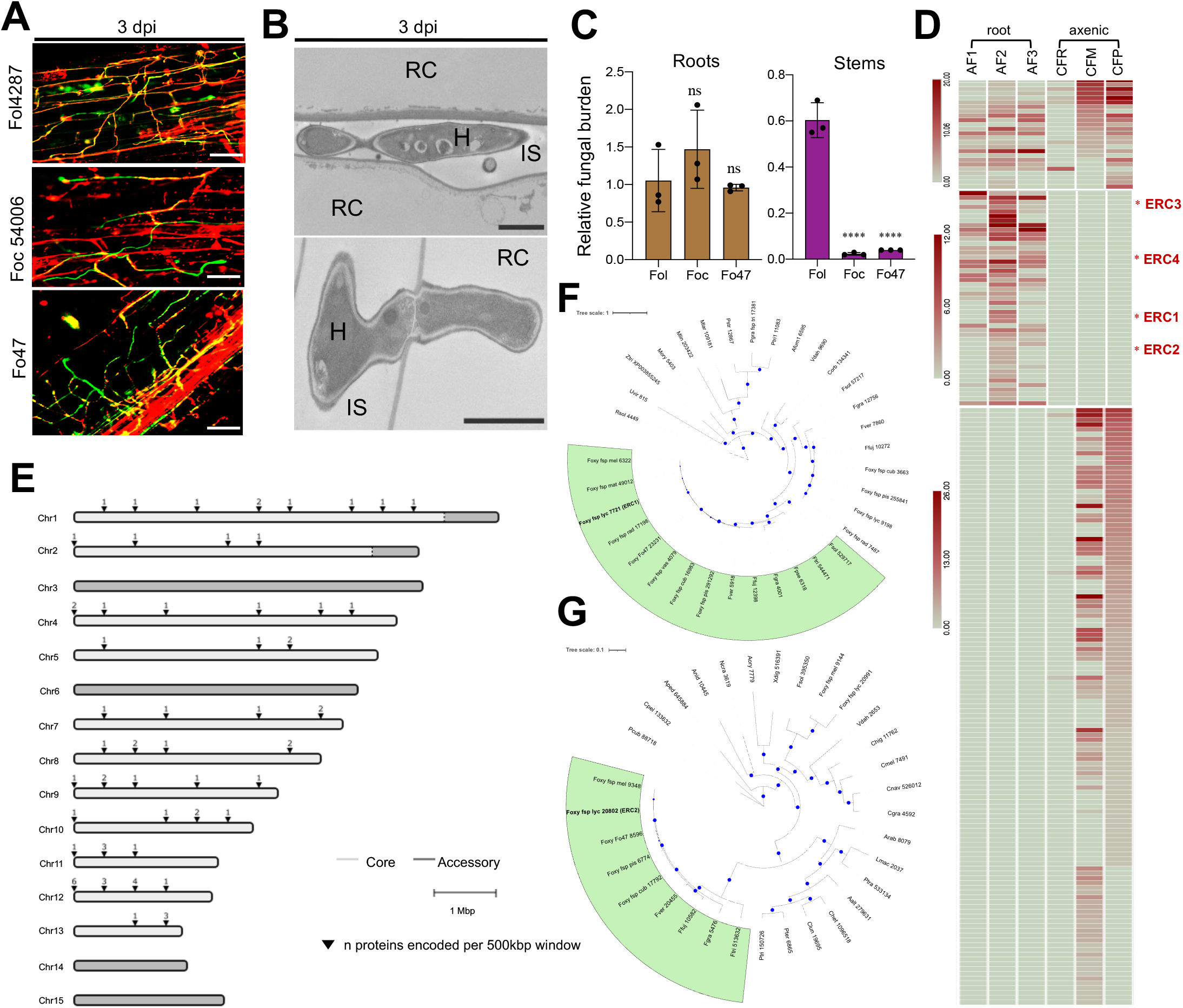
*F. oxysporum* secretes a suite of conserved core effectors in the root intercellular space. (**A**) Confocal microscopy of tomato root colonization of *F. oxysporum* isolates Fol4287 (tomato pathogen), Foc54006 (banana pathogen) or Fo47 (endophyte) expressing 3xclover at 3 dpi. Fungal fluorescence (mClover3-green) is overlaid with propidium iodide staining of plant cell walls (red). Scale bars, 25 µm. (**B**) TEM micrographs showing hyphae of Fol4287 (H) growing intercellularly (top) or penetrating a root cell (RC; bottom). IS, intercellular space. Scale bars, 2 µm (top); 1 µm (bottom). (**C**) Fungal burden in roots and stems of tomato plants inoculated with the indicated Fo isolates were measured by real time (qRT)-PCR of the Fo *actin* gene using total DNA extracted at 12 dpi. Fo DNA was calculated using the threshold cycle (ΔΔCt) method, normalized to the tomato *gadph* gene. Error bars indicate standard deviation (s.d.); n = 3 biological replicates. Asterisks indicate statistical significance versus Fol4287 (one way ANOVA, Bonferroni’s multiple comparison test, p<0.05). ns = non-significant. Experiments were performed three times with similar results. (**D**) Heat map showing absolute counts of unique peptides of Fol4287 identified by LC-MS/MS in three independent samples of tomato root apoplastic fluid (AF) at 3dpi (AF1,2,3) or in a single sample of filtrate from axenic cultures in minimal medium with tomato crushed roots (CFR) or sucrose as carbon source (CFM) or potato dextrose broth (CFP). Putative effectors ERC1-4 are highlighted with a red asterisk. Note differences in scale between sections of the heat map. (**E**) Chromosomal distribution plot of the genes encoding *F. oxysporum* Fol4287 proteins identified in tomato AF, showing their localization exclusively in core genomic regions. Core and LS regions are shown in light and dark grey, respectively. (**F** and **G**) Maximum likelihood phylogenetic trees based on the aligned amino acid sequences of the FOXG_11583 (ERC1) (**F**) and FOXG_04534 (ERC2) proteins (**G**). Number indicates MycoCosm protein ID. Size of blue dots represents bootstrap support for the branch with maximum 100 bootstraps. Fungal species included in the analysis are listed in table S4.

### *F. oxysporum* secretes a battery of core effector proteins in the root intercellular space

During plant infection, pathogenic and mutualistic fungi secrete effectors to modulate host responses (2). Some effectors are highly specific for a given pathogen species while others are conserved across a broad range of fungal pathogens and endophytes (26). Given the predominantly apoplastic growth of Fo during the early biotrophic stage (Fig.1, A and B), we performed discovery proteomics of tomato root apoplastic fluid (AF) collected at 3 dpi to search for potential effectors. Liquid chromatography-mass spectrometry (LC-MS) analysis identified a total of 72 Fol4287 proteins present in AF, 32 of which were consistently detected across three biological replicates (Fig. 1D, fig. S1C and D, Data S1). Twenty-five of the 72 apoplastic fungal proteins were also found in the filtrates of axenically grown fungal cultures, while 47 were only detected *in planta* (Data S1). The latter include different classes of cell wall degrading enzymes such as polygalacturonase, pectate lyase, glucanase or galactosidase, as well as other enzymes such as a choline dehydrogenase, a copper amine oxidase or a carbonic anhydrase (CA). CAs were previously associated with pH adaptation, CO_2_-sensing and virulence of fungal pathogens of humans such as *Cryptococcus neoformans* (27). Moreover, plant CAs were recently shown to regulate basal immune responses in tomato (28). Interestingly, while CAs from human fungal pathogens belong to the beta-CA class and lack a secretion signal (29), the Fo CA identified in AF belongs to the poorly characterized alpha-CA class and carries a signal peptide, suggesting a possible adaptation to the plant pathogenic lifestyle.

In an attempt to identify *bona fide* effector candidates we looked for proteins that 1) were specifically present in AF; 2) were consistently detected in all three biological AF replicates; 3) contain a predicted N-terminal secretion signal peptide; 4) lack predicted transmembrane domains; 5) contain multiple cysteines. Four of the identified Fo proteins fulfilling all five criteria were selected for further analysis and named <u>E</u>arly <u>R</u>oot <u>C</u>ompatibility (ERCs) effectors (Fig. 1D, fig. S1E and F, Data S1). ERC1 (FOXG_11583) carries a putative cellulose-binding domain (30), while ERC3 (FOXG_16902) has a α-L-arabinofuranosidase domain similar to those reported in effectors from the rice blast pathogen *Pyricularia oryzae* or the corn smut fungus *Ustilago maydis* (31, 32). Intriguingly, ERC2 (FOXG_04534) and ERC4 (FOXG_08211) both carry a predicted lytic polysaccharide monooxygenase (LPMO) domain. LPMOs cleave cellulose, chitin and other polysaccharides through a novel oxidative mechanism and have been suggested to act on crystalline surface regions of the substrate to create attachment sites and enhance accessibility for canonical glycoside hydrolases (33). LPMOs have been associated with plant pathogenicity, and pectin-cleaving LPMOs were recently shown to drive plant infection in the Irish potato famine oomycete pathogen *Phytopthora infestans* (34, 35). Thus, the four putative Fol4287 effectors detected in AF are proteins associated either with binding or modification of plant cell walls. This is in line with a recent study that identified secreted fungal proteins related with cell wall modification as genetic determinants of endophytism in phylogenetically distant members of the *A. thaliana* root mycobiome (36).

We noted that almost all Fo proteins secreted in root AF, including ERC1-4, are encoded on core genomic regions that are shared across the entire Fo species complex (Fig. 1E). This contrasts sharply with the previously reported SIX effector genes, which are located on lineage-specific regions (14, 15). Moreover, in contrast to most SIX effectors, ERCs have predicted homologues outside Fo, including many asco- and basidiomycete species with distinct lifestyles including both phytopathogens and non-pathogens (Fig.1F and G; fig. S1G, table S4, Data S2 to S4). Interestingly, ERC1 and ERC2 each have a paralog in Fol4287 (FOXG_12855 and FOXG_18882, respectively), which cluster separately and are also conserved across the entire Fo species complex, suggesting the presence of functionally diverged members of the ERC1 and ERC2 effector families (Fig.1 F and G). FOXG_18882 shows 56% identity to ERC2, but is much shorter (97 aa versus 253aa) and clusters with predicted homologs from *F. solani* and Fo f. sp. *melonis* (Fig. 1G). ERC3 and ERC4 homologs were also found across ascomycetes and some basidiomycete groups such as the *Agaricomycotina* and *Ustilaginomycotina*, but not in the *Pucciniomycotina* (fig. S1G, Data S4 and S5). We conclude that Fo4287 secretes an array of putative core effectors during early stages of tomato root colonization, that are conserved in diverse fungi including both pathogenic and non-pathogenic plant-associated species.

### ERCs are upregulated during the early biotrophic infection stage

A hallmark of hemibiotrophic pathogens is a switch from biotrophy to necrotrophy coinciding with a major shift in the gene expression pattern (37). In contrast to air-borne plant pathogens, the transcriptional dynamics of hemibiotrophic fungal root infection has not been explored in detail (38). Here we performed RNA sequencing (RNAseq) of tomato roots inoculated with Fol4287 during early stages of biotrophic growth in the root cortex (1, 2 and 3 dpi) as well as during colonization of the root vascular tissue (7 dpi), which marks the transition from biotrophy to necrotrophy. Among the genes encoding proteins previously identified in AF (Data S1), 36 (77%) showed a marked transcriptional upregulation during early infection stages as compared to axenic growth conditions (fig. S2A). Analysis of differentially expressed genes (DEGs; log2 fold change >2, P <0.05) and Principal Component Analysis revealed a major shift in the transcriptional profile, both between axenic and *in planta* conditions (Fig. 2A and Data S6 to S9) as well as between early (1, 2, 3 dpi) and late (7 dpi) stages of infection (fig. S2B). The total number of DEGs ranged from 1500-1800 upregulated and 600-800 downregulated genes depending on the time point (Fig. 2A). Gene ontology (GO) enrichment analysis showed an abundance of fungal transcripts involved in amino acid biosynthesis and metabolic processes during the early colonization stages (fig. S2C, Data S11), which is indicative of biotrophic growth as previously shown for the interaction between *Piriformospora indica* and *Arabidopsis* (39).

**Fig. 2.**
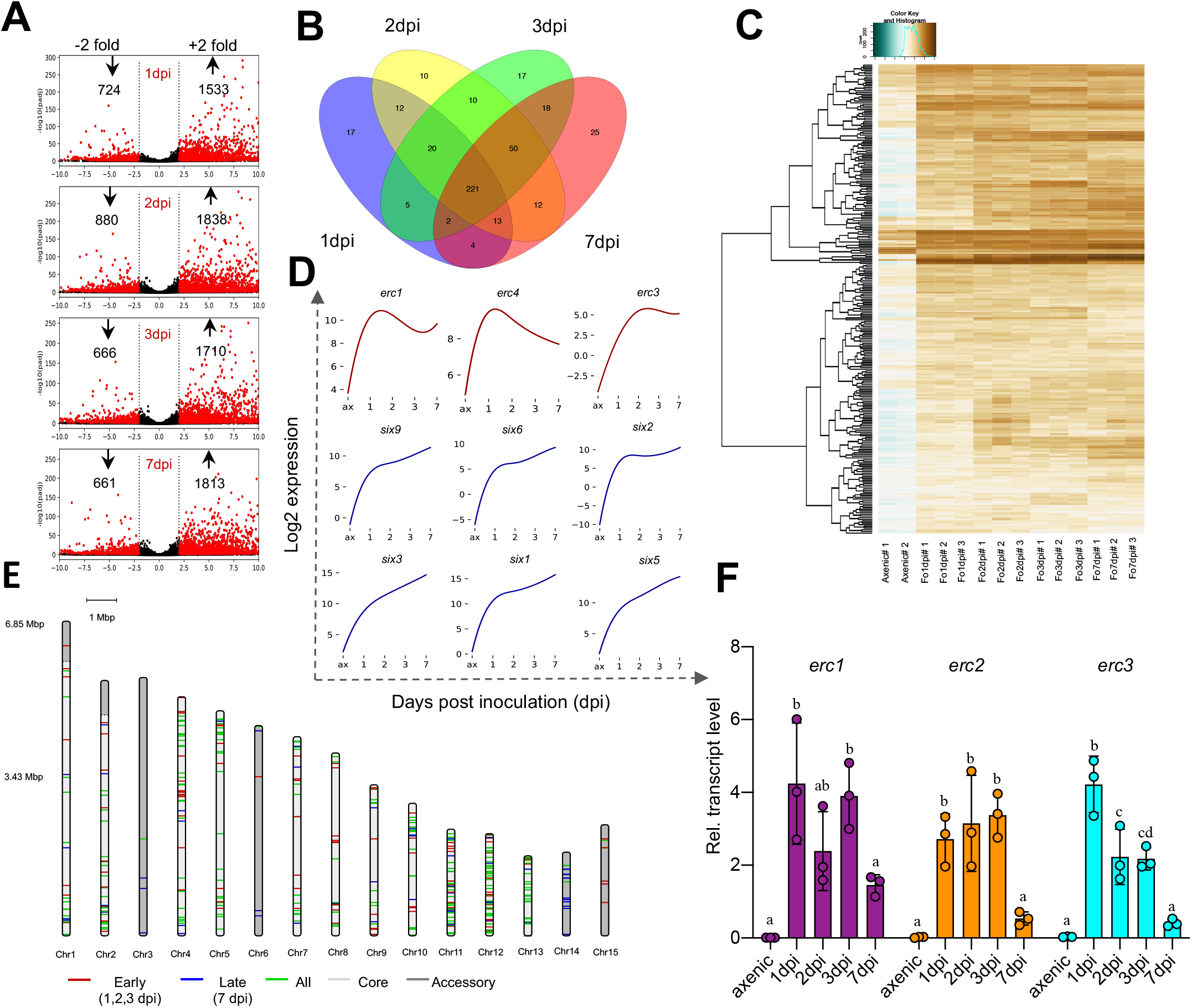
ERCs are upregulated during early biotrophic stages of infection. (**A**) Volcano plot showing pairwise differential expression analysis of Fol4287 genes at the indicated time points after inoculation in tomato roots versus axenic culture. Significantly differentially expressed genes are in red. (**B**) Venn diagram of Fol4287 genes encoding predicted secreted proteins (SPs) upregulated in tomato roots at the indicated time points after inoculation versus axenic culture. A core set of 221 SPs upregulated *in planta* were identified across all the analysed timepoints. (**C**) Hierarchical clustering and heatmap showing log_2_-fold changes of Fol4287 genes encoding secreted proteins differentially expressed in tomato roots versus axenic culture (*t* test, *P <* 0.05). Brown, upregulated genes; blue, downregulated genes. (**D**) Representative expression profiles of the indicated candidate effector genes plotted as log2 expression versus days post inoculation (dpi). Profiles in red correspond to the indicated ERC genes located on core genomic regions, showing upregulation at early time points with a drop at 7dpi. Profiles in blue correspond to known SIX effector genes encoded on the lineage specific (LS) chromosome 14 showing maximum upregulation at 7dpi. (**E**) Chromosomal distribution of *in planta* upregulated Fol4287 genes encoding predicted secreted proteins. Red bands are genes preferentially expressed at early infection stages (early to late expression ratio > 0.25). Blue bands are genes preferentially expressed at late infection stages (early to late expression ratio < 0.25). Green bands are genes expressed at both early and late infection stages (differentially expressed in at least one early plus the 7dpi dataset versus axenic). Early refers to 1-3dpi; late refers to 7dpi. Core and LS genomic regions are shown in light and dark grey, respectively. (**F**) Relative transcript levels of genes *FOXG_11583* (*erc1*), *FOXG_04534* (*erc2*) and *FOXG_16902* (*erc3*) were measured by qRT-PCR of cDNA obtained from Fol4287 grown in minimal medium (axenic) or from roots of tomato plants inoculated with Fol4287 at 1, 2, 3 or 7 dpi. Transcript levels were calculated using the threshold cycle (ΔΔCt) method and normalized to the Fol4287 peptidyl prolyl isomerase (*ppi*) gene. Error bars indicate standard deviation (s.d.); n = 3 biological replicates. Different letters indicate statistically significant differences according to one way ANOVA, Bonferroni’s multiple comparison test (p < 0.05).

Around 6% (436) of the 6894 *in planta*-induced genes during the early colonization timepoints tested encode predicted secreted proteins. While many of these display fluctuations in the expression profile across the different infection stages, we also found a set of 221 genes that were significantly upregulated at all infection timepoints tested (Fig. 2B). Importantly, among the *in planta* expressed genes encoding predicted secreted proteins are previously characterized Fol4287 effectors belonging to the metalloprotease and serine protease families (40) (fig. S2D). Hierarchical clustering and expression analysis revealed two distinct patterns of expression for *in planta* upregulated genes encoding secreted proteins: 1) those showing maximum transcript levels during the early stage (1, 2 and 3 dpi) and a subsequent drop at 7 dpi, and 2) those showing a progressive induction with an expression peak at 7 dpi (Fig. 2C and D). The first category includes the *ERC* genes, among others, while the second category includes the *SIX* genes. Strikingly, most of the early upregulated genes are located on core genomic regions (except some genes located on LS chromosome 15), while the late upregulated effector genes are predominantly located on LS regions, particularly on the so-called pathogenicity chromosome 14 (Fig. 2E).

Regarding the expression of *ERC* genes, the results from RNAseq are generally in line with those from proteomics of AF at 3 dpi (Fig. 1D and E). We further performed qRT-PCR analysis of genes *erc1, 2, 3* and *4* confirmed their upregulation *in planta* during the early infection stage and the subsequent downregulation of three of the genes at 7 dpi (Fig. 2F, fig. S2E). Taken together, our results suggest that Fol4287 undergoes a major transcriptional shift between the early stage of biotrophic root colonization (1, 2, 3 dpi) and the later stage at 7 dpi coinciding with the onset of growth in the xylem. The finding that core effectors such as ERCs are specifically upregulated during the initial stage suggests a potential role in the early establishment of fungus-plant compatibility.

### ERCs contribute to host plant infection and suppression of root immunity

To test the role of ERCs in host plant colonization and virulence, we generated isogenic Fol4287 deletion mutants lacking either the *FOXG_11583, FOXG_04534, FOXG_16902* or *FOXG_08211* gene, which were named Δ*erc1*, Δ*erc2*, Δ*erc3* and Δ*erc4* respectively (fig. S3A to S3E). Phenotypic analysis of these mutants revealed no detectable effect on vegetative growth, colony morphology or stress resistance on different media (fig. S3G). However, the Δ*erc1*, Δ*erc2* and Δ*erc3* mutants, caused significantly reduced mortality on tomato plants and accumulated less *in planta* fungal biomass than the Fol4287 wild type strain (Fig. 3A to C and fig. S3F, S3H and S3I). Moreover, trypan blue staining confirmed that these mutants were less efficient in colonizing tomato roots than the wild type (fig. S3J). Importantly, reintroduction of the wild type allele (fig. S3K) fully restored virulence and root colonization in the Δ*erc1*, Δ*erc2* and Δ*erc3* mutants (Fig. 3 A to C, fig. S3H and S3I).

**Fig. 3.**
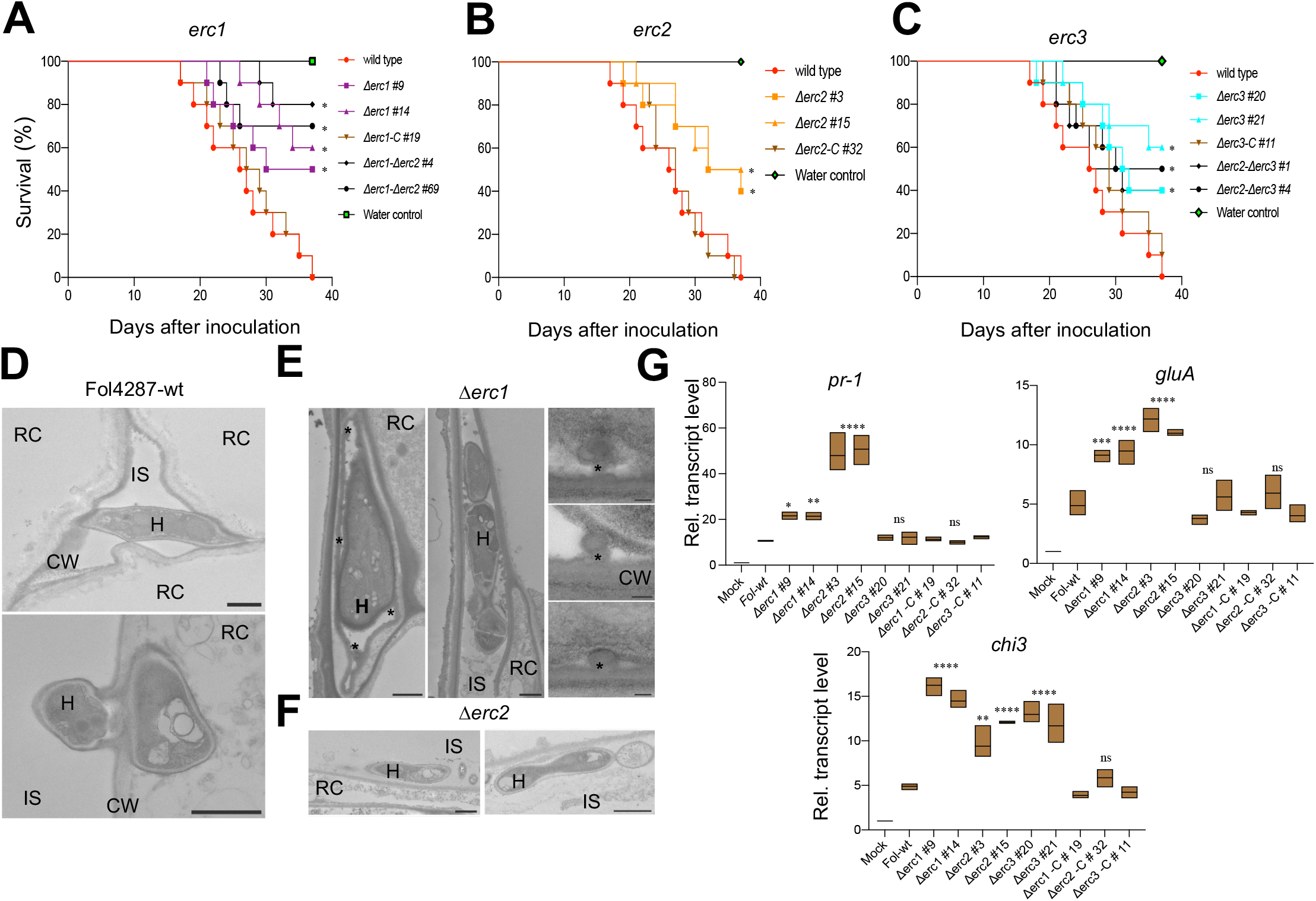
ERCs contribute to root colonization, virulence and suppression of the plant immune responses. (**A** to **C**) ERCs have a role in virulence of *F. oxysporum*. Kaplan-Meier plot showing the survival of tomato plants inoculated with the Fol4287 wild type strain or the indicated single or double gene knockout mutants. Number of independent experiments = 3; 10 plants/treatment. Data shown are from one representative experiment. \**P* <0.05, versus the wild type according to log-rank test. Note that mortality caused by the *Δerc1, Δerc2* and *Δerc3* single mutants is significantly lower than that caused by the wild type strain while mortality caused by the *Δerc1Δerc2 and* Δ*erc2Δerc3* double mutants is not significantly different from that of the respective single mutants. (**D**) TEM micrographs showing hyphae (H) of the Fol4287 wild type (wt) growing between (top) or penetrating into tomato root cells (RC; bottom). CW; plant cell wall, IS; intercellular space. Scale bars, 1 µm. (**E** and **F**) TEM micrographs showing hyphae (H) of the Fol4287 Δ*erc1* (e) or Δ*erc2* (f) mutants growing between tomato root cells (RC). Note in (**E**) that hyphae are encapsulated by protrusions of amorphous granular material (asterisks) and in (**F**) hyphae are located close to the root periphery. Scale bars in (**E**), 1 µm in left and centre images; 0.1 µm in right image; in (**F**) 1 µm. (**G**) Transcript levels of tomato defence genes *pr-1, gluA* and *chi3* were measured by RT-qPCR of cDNA obtained from tomato roots at 2 days after inoculation with the indicated fungal strains. Transcript levels were calculated using the ΔΔCt method, normalized to the tomato *gadph* gene and expressed relative to those of the uninoculated control (H_2_O). Asterisks indicate statistically significant differences according to one way ANOVA, Bonferroni’s multiple comparison test (p < 0.05). Box length indicates s.d.; n = 3 biological replicates. Experiments were performed three times with similar results.

TEM analysis of tomato roots inoculated with the Δ*erc1* or Δ*erc2* mutants at 3 dpi revealed a lack of penetration of root cortical cells (Fig. 3E and 3F, fig. S4B) as compared to the wild type strain where successful penetration events were frequently observed (Fig. 1B and 3D, fig. S4A). Moreover, in the Δ*erc1-*tomato interaction we noted the presence of characteristic protrusions from the cell walls of root cortex cells that encapsulated the fungal hyphae, as well as the secretion of an amorphous granular material (asterisks in Fig. 3E, arrowheads in fig. S4B). Similar structures have been reported previously in plants infected with vascular fungal pathogens such as *Fusarium, Verticillium* or *Ceratocystis* and were suggested to contribute to plant resistance by preventing plant cell wall degradation and inhibiting hyphal spread (41-43). In line with this idea, in contrast to the wild type strain which was detected inside the vascular bundles and parenchyma cells, the Δ*erc1* and Δ*erc2* mutants remained largely restricted to intercellular growth in the root cortex (Fig. 3E and 3F).

The distinct ultrastructural phenotypes of the Δ*erc* single mutants suggest that these effector proteins may have different, non-redundant roles in promoting host root colonization. To test this idea, we generated Δ*erc1*Δ*erc2* and Δ*erc2*Δ*erc3* double knockout mutants. Interestingly, the Δ*erc1*Δ*erc2* strains caused slightly lower mortality compared to the wild type and Δ*erc* single mutants (Fig. 3A) and was further reduced in plant colonization as determined by trypan blue staining and qRT-PCR (fig. S4C to S4E). By contrast, the Δ*erc2*Δ*erc3* double mutants displayed similar levels of virulence levels and fungal burden as the single mutants (Fig. 3C and fig. S4C to S4E). To test whether ERCs contribute to suppression of host defence, we measured the transcript levels of known plant defence genes at 2 dpi. We found a 2 to 4 fold increase in upregulation of the *pr-1, gluA* and *chi3* genes encoding pathogenesis-related protein 1, basic β-1,3-glucanase and acidic chitinase, respectively (12), in tomato roots infected with the Δ*erc1*, Δ*erc2 and* Δ*erc3* mutants compared to those infected with the wild type or the complemented strains (Fig. 3G). Taken together, these results suggest that ERCs contribute to host plant infection by Fol and may contribute to suppression of the plant immune response.

### ERCs are also required for endophytic colonization and biocontrol activity of a non-pathogenic Fo strain

It has been known for decades that Fo can grow as an endophyte on roots of non-host plants without inducing detectable disease symptoms (22, 23). Moreover, certain Fo isolates appear to be non-pathogenic, such as Fo47, a well-characterized biocontrol strain that was originally isolated from a soil naturally suppressive to *Fusarium* wilt (24). Fo47 triggers endophyte-mediated resistance (EMR) against plant pathogenic forms of Fo (44). Here we confirmed that co-inoculation of tomato plants with Fo47 and Fol4287 resulted in a marked reduction of mortality caused by Fol4287, as well as in a decrease of *in planta* Fol4287 biomass, when compared to plants inoculated with Fol4287 alone (fig. S5A to S5C). Although the hyphae of both Fol4287 and Fo47 were able to penetrate the root endodermis cells and grow into xylem vessels, Fol4287 showed more profuse spread compared to Fo47 (Fig. 4A). Interestingly, TEM analysis revealed the presence of fungal membrane tubules in penetration hyphae of Fol4287 (Fig. 4A, fig. S5D and S5E). Similar structures were previously reported during invasive growth of plant symbionts and pathogens (45). On the other hand, we frequently observed aborted penetration events of tomato cell by Fo47 hyphae that were devoid of cytosol (Fig. 4A, arrows; fig. S5F to S5H). In the rare occasions where Fo47 hyphae were observed inside xylem vessels, they were often encapsulated by an amorphous granular material which also encrusted the plant cell walls and blocked the pits between the xylem vessel and adjacent cells (Fig. 4A asterisks and fig. S5I arrowheads). By contrast, such material was rarely observed in xylem vessels colonized by Fol4287, where hyphae successfully spread between xylem vessels (Fig 4A arrowheads and fig. S5J), and was completely absent from the xylem pits of uninoculated plants (fig. S5K and S5L). The deposition of amorphous granular material, as well as phenolic infusion, lignification or incorporation of calcium into pit membranes was previously suggested to contribute to the defence against vascular wilt pathogens such as Fol, *Verticillium albo-atrum* (41) or *Ceratocystis fimbriata*, by inhibiting the spread of pathogen hyphae and preventing plant cell wall degradation (41-43). Overall, our findings indicate that the commitment of Fo towards a pathogenic or endophytic lifestyle likely occurs at the level of endodermis penetration and vascular colonization which is successfully completed by the pathogenic strain Fol4287, but are either inhibited or blocked in the endophytic isolate Fo47. Our results also confirm that the biocontrol strain Fo47 is able to colonize tomato roots to a certain extent and to protect the plant against the pathogenic form Fol4287.

**Fig. 4.**
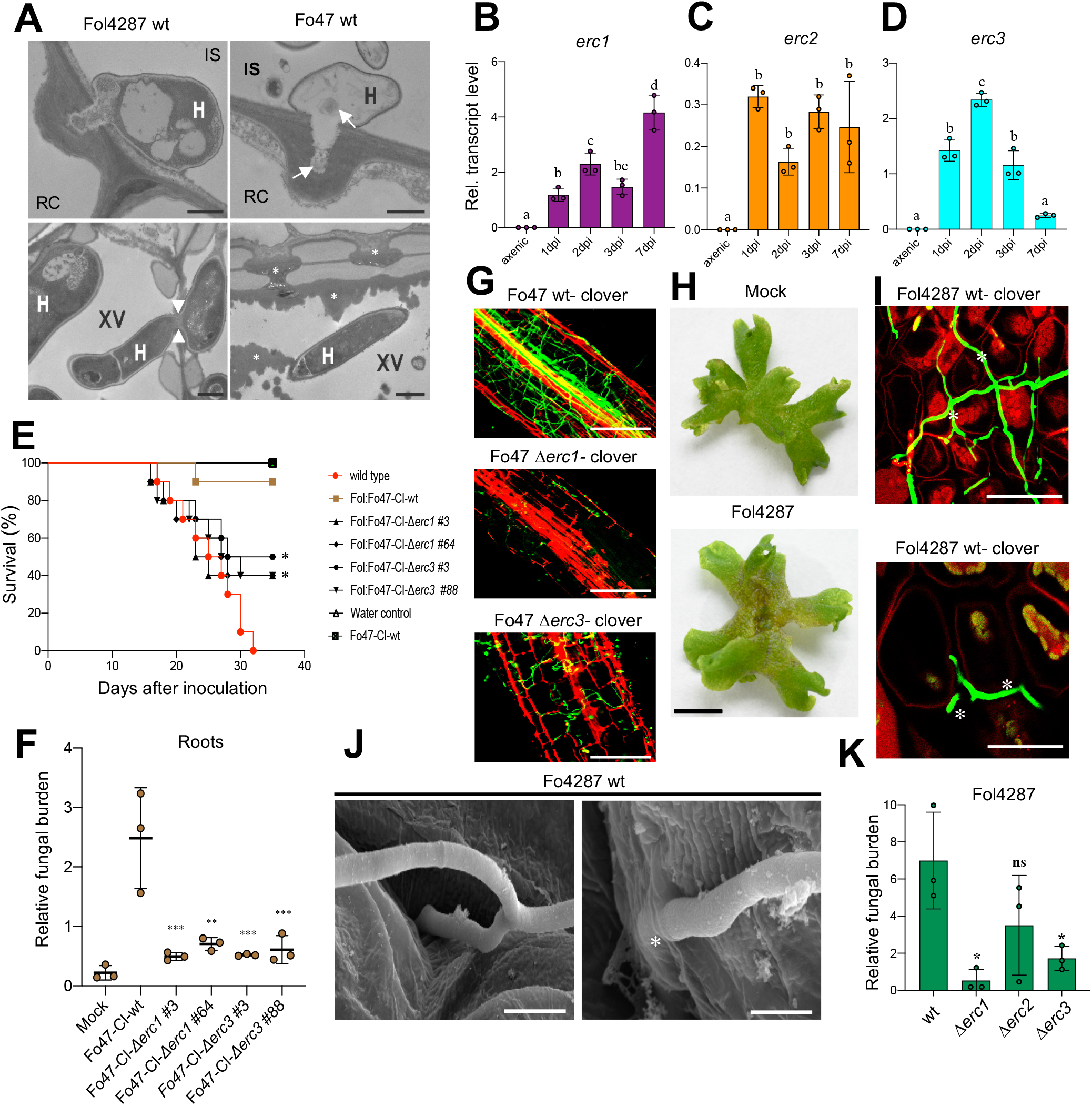
ERCs evolved as fungal compatibility factors on both host and non-host plants. (**A**) TEM micrographs showing hyphae (H) of Fol4287 and Fo47 attempting penetration of tomato root cells (RC) (top) or growing inside xylem vessels (XV; bottom). Note that the penetration hypha of Fo47 is devoid of cytosol and contains only remnants of cell components (arrows). Arrowheads indicate a Fol4287 hypha penetrating an adjacent xylem vessel. Asterisks indicate the deposition of amorphous granular material encapsulating a Fo47 hypha in a xylem vessel or blocking the pits between vessels to inhibit cell-to-cell movement of the fungus. CW, cell wall; IS, intercellular space. Scale bars= 1 µm in top images; 2 µm in bottom images. (**B** to **D**) Relative transcript levels of the genes *FOZG_11686* (*erc1*), *FOZG_02496* (*erc2*) and *FOZG_12886* (*erc3)* in isolate Fo47 (biocontrol strain) were measured by qRT-PCR of cDNA obtained from fungal mycelium grown in minimal medium (axenic) or from roots of tomato plants inoculated with Fo47 at 1, 2, 3 or 7 dpi. Transcript levels were calculated using the threshold cycle (ΔΔCt) method and normalized to the Fo peptidyl prolyl isomerase gene *(ppi)*. Error bars indicate standard deviation (s.d.); n = 3 biological replicates. Different letters indicate statistically significant differences according to one way ANOVA, Bonferroni’s multiple comparison test (p < 0.05). (**E**) Kaplan-Meier plot showing the survival of tomato plants inoculated with the Fol4287 or Fo47-mClover wild type strains alone or co-inoculated with Fol4287 plus Fo47-mClover wild type or the indicated Fo47-mClover Δ*erc* mutants. Number of independent experiments = 3; 10 plants/treatment. Data shown are from one representative experiment. \**P* <0.05, versus Fol4287 alone according to log-rank test. Note that protection provided by the Fo47-mClover *Δerc1*, and *Δerc3* mutants is significantly lower than that provided by the Fo47-mClover wild type strain. (**F**) Fungal burden in roots of tomato plants inoculated with the indicated Fo isolates was measured by qRT-PCR of the Fo47 specific gene (*FOBG_10856*) using total DNA extracted at 10 dpi. Fo DNA was calculated using the threshold cycle (ΔΔCt) method, normalized to the tomato *gadph* gene and expressed relative to that in roots inoculated with Fo47-mClover wt. Error bars indicate standard deviation (s.d.); n = 3 biological replicates. Asterisks indicate statistical significance versus Fo47-mClover wild type (one way ANOVA, Bonferroni’s multiple comparison test, p<0.05). The experiment was performed three times with similar results. (**G**) Confocal microscopy images showing tomato roots inoculated with the Fo47-mClover wild type strain or the indicated *Fo47-*mClover Δ*erc* mutants at 3 dpi. Fungal fluorescence (green) is overlaid with propidium iodide staining of plant cells (red). Scale bars, 25 µm. (**H**) Macroscopic disease symptoms of *M. polymorpha* Tak-1 thalli 5 days after dip inoculation with 10^5^ microconidia ml^-1^ of Fol4287 or water (mock). Images are representative of three independent experiments. Scale bar, 1 cm. (**I**) Confocal microscopy showing intercellular growth of Fol4287-mClover on a Tak-1 thallus at 3 dpi. Fungal fluorescence (green) is overlaid with propidium iodide staining of plant cells (red). Asterisks indicate intercellular hyphal growth. Scale bar, 25 µm. (**J**) SEM micrographs showing hyphae (H) entering thalli of *Marchantia*, either intercellularly (arrow) or by direct penetration (asterisks). Scale bars= 5 µm in top image; 2 µm bottom image. (**K**) Fungal burden in 3-week-old *Marchantia* thalli inoculated with the Fol4287 wild type strain or the indicated Δ*erc* mutants was measured by qRT-PCR of the Fo *actin* gene using total DNA extracted at 6 dpi. The relative amount of fungal DNA was normalized to the *Mp* EF1a gene. Statistical significance versus wt (p < 0.05, Unpaired t-test) is indicated by an asterisk. ns = non-significant. Error bars indicate s.d. (n = 3).

Because ERCs are conserved across the Fo species complex including non-pathogenic forms such as Fo47, we reasoned that they might contribute to pathogenic as well as endophytic root colonization. In support of this, *erc1, erc2* and *erc3* were transcriptionally upregulated during early stages of tomato root colonization in the non-pathogenic isolate Fo47 (biocontrol interaction) or the banana pathogenic isolate Foc54006 (non-host interaction), as previously observed in Fol4287 (Fig. 4B to 4D and fig. S6A to S6C). Moreover, we found that the *erc1* and *erc2* orthologs of the vascular wilt pathogen *Verticillium dahliae* VdLS17 (46) (VDAG_04446 and VDAG_07135) were also transcriptionally induced during early stages of tomato root colonization (fig. S6D to S6F).

We next asked whether ERCs contribute to root colonization and biocontrol activity of the non-pathogenic isolate Fo47. Indeed, Δ*erc1* and Δ*erc3* deletion mutants obtained in a Fo47 mClover3 background were reduced in their ability to colonize tomato roots compared to the wild type Fo47 mClover3 strain (Fig. 4G). Importantly, these mutants were also significantly less efficient in colonizing and protecting tomato plants against mortality caused by Fol4287 and in reducing wilting due to the pathogenic isolate (Fig. 4E and 4F, fig. S5C). Overall, these results suggest that ERCs are used by different Fo isolates with contrasting lifestyles (pathogenic and endophytic) during early infection stages to establish associations with plant roots.

### ERCs contribute to Fo infection on the non-vascular land plant *Marchantia polymorpha*

The broad distribution of ERC homologues across the fungal kingdom suggests a conserved role of these core effectors in plant colonization. To experimentally test this idea, we took advantage of a recently established Fo infection model in the liverwort *Marchantia polymorpha* (47), which has emerged as a non-vascular model for molecular plant-microbe interactions (48). As previously reported (47), we found that Fol4287 causes visible disease syptoms on *Marchantia* thalli and displays mostly intercellular hyphal growth similar to that observed in the cortex of tomato roots (Fig 4H and 4I). Scanning electron microscopy showed Fol4287 hyphae entering the thalli intercellularly, either by growing between cells or through air pores (Fig. 4J and fig. S7A). We also observed occasional events of direct penetration (Fig. 4J). Similar to infection of tomato plants, transcript levels of the *erc1, erc2* and *erc3* genes were markedly upregulated during growth of Fol4287 in *Marchantia* compared to the axenic control (fig. S7B to S7D). Interestingly, *Marchantia* thalli inoculated with the Fol4287 Δ*erc1*, Δ*erc2 or* Δ*erc3* mutants showed a slight reduction in the severity of disease symptoms compared to those inoculated with the wild type strain (fig. S7E). Moreover, the fungal burden in the thalli inoculated with the Δ*erc* mutants at 6 dpi was lower than in those inoculated with the Fol4287 wild type strain, although the difference was only significant for the Δ*erc1*and Δ*erc3* mutants (Fig 4K). We conclude that ERCs, particularly ERC1 and ERC3, contribute to infection of Fol4287 on the non-vascular bryophyte model *M. polymorpha*.

Besides causing disease in plants, *F. oxysporum* has also been reported as an opportunistic pathogen of humans (49). Fol4287 was previously shown to cause mortality on immunodepressed mice and larvae of the invertebrate insect model host *Galleria mellonella* (50, 51). Here we found that mortality caused by the Δ*erc1*, Δ*erc2* or Δ*erc3* mutants on *G. mellonella* larvae did not differ significantly from that caused by the Fol4287 wild type strain, suggesting that ERCs are dispensable for infection on animal hosts (Fig. S7F to S7H). Collectively, this data suggests that ERC effectors contribute to fungal infection on evolutionarily distant plant host lineages independent of the presence of a true vasculature.

## Discussion

The soilborne vascular pathogen Fo causes systemic infections and wilting on a broad range of crops, with individual strains exhibiting exquisite host specificity determined by LS effector proteins (7). However, pathogenic isolates can also colonize roots of non-host plants without causing wilt and hence behave as endophytes (7). Such multi-host colonization must likely involve pathogen factors that are conserved across a wide range of Fo isolates, but the nature of these molecules has so far remained elusive. By integrating discovery proteomics with early-stage RNA-seq and targeted gene knockout analysis, we show here that colonization of multiple plant hosts by Fo is mediated by a set of conserved compatibility factors termed <u>E</u>arly <u>R</u>oot <u>C</u>ompatibility effectors (ERCs), which are encoded by core genomic regions and secreted into the root apoplast during the initial stages of infection.

Plant cell wall modification and modulation of apoplastic immunity appear to be essential for Fo establishment, as suggested by the large proportion of the apoplastic fungal proteome representing cell-wall modifying enzymes and small secreted proteins. A similar enrichment of cell wall degrading enzymes was also observed in the apoplastic fluid of rice leaves infected by the rice-blast pathogen *P. oryzae* or during symbiotic root interactions with the endophytic fungus *Piriformospora indica* (52, 53). Collectively, these findings suggest that cell wall modification and suppression of host immunity represent conserved fungal mechanisms for plant colonization.

Our transcriptomic datasets from early infection stages (1-3 dpi) identified ∼230 predicted secreted proteins. Interestingly, most of these early-induced effector candidates are encoded on core-genomic regions, whereas most of the late-induced genes (7dpi) are encoded on LS chromosome 14. This suggests that pathogenic lifestyle transitions in Fo are accompanied by transcriptional regulation of different waves of effectors as previously reported in the hemibiotrophic fungal pathogen *Colletotrichum* (37). Among the secreted proteins overlapping in the proteomic and transcriptomic datasets, we identified four early effector candidates which are endoded on core genomic regions and conserved across the Fo species complex as well as other fungi. These ERCs are upregulated during colonization of both host and non-host plants independently of the fungal lifestyle, indicating that they have a broadly conserved role in fungus-root associations rather than a host-specific role in vascular colonization and wilting, as described previously for the LS-encoded SIX effectors.

Targeted deletion of *ERC* genes in the tomato pathogenic isolate Fol4287 resulted in impaired root colonization, reduced virulence and rapid activation of plant immune responses, while in the non-pathogenic isolate Fo47 it led to a reduction in root colonization and biocontrol ability against the pathogenic Fo isolate. It is worth noting that all four ERC effectors contain predicted domains associated either with binding or modification of plant cell walls. This suggests that these effectors may either function in the mobilization of carbohydrates in the apoplast or in shielding the fungus from plant immune responses. TEM analysis of the Δ*erc1* and Δ*erc2* mutants support this hypothesis as both mutants are unable to transition from inter- to intracellular growth and are blocked in the intercellular space through distinctive plant cell wall protrusions. Interestingly, a large-scale genomic analysis across diverse fungal genera in *A. thaliana* roots identified a set of proteins with cell wall modifying function as genetic determinants of endophytism (36). The precise molecular role of ERCs is currently unclear. Intriguingly, a recent work in *P. infestans* showed that a secreted pectin mono-oxygenase with a LPMO domain contributes to plant infection by cleavage of pectin (35). We speculate that LPMO effectors such as ERC2 might be used by Fo to dampen recognition by the host and to cleave the polysaccharide backbone of the plant cell wall to aid pathogen accommodation.

We have a limited understanding on how plants engage with beneficial microbes and at the same time restrict pathogens, most likely with an immune thermostat to select for nature of the interaction (54). Recent evidence suggests that plant roots employ cell-layer-specific programs to respond to beneficial and pathogenic microbes (55). Our comparative TEM analysis of the pathogenic isolate Fol4287 and the biocontrol isolate Fo47 revealed significant differences in the ultrastructural development of the two isolates in the inner root cell layers, where growth of the Fo47 hyphae was abrogated while early growth in root cortical cells was similar between the two strains. This is in line with the finding that ERC genes are expressed predominantly during the initial infection stage and thus are likely to play a role in early root colonization which is conserved between the two fungal isolates. By contrast, host-specific LS effectors expressed at later infection stages determine entry and colonization of the vasculature and the switch to pathogenicity. Importantly, our results in the non-vascular bryophyte model *M. polymorpha* confirm that ERCs are induced during plant intercellular growth of Fo independent of vasculature signatures. The finding that some ERCs also contribute to Fo infection on this evolutionarily distant plant lineage further supports a broadly conserved role of ERCs in fungus-plant interactions.

Taken together, our study uncovers a suite of previously uncharacterized early root compatibility effectors, which are secreted by the vascular wilt fungus Fo during the initial asymptomatic infection stages. ERCs contribute to plant colonization by both pathogenic and non-pathogenic Fo isolates on a wide range of host and non-host plants, ranging from tomato to the liverwort *M. polymorpha* which lacks a differentiated vasculature. While the modes of action of ERCs are currently unknown, our results suggest that ERCs may target evolutionarily ancient plant processes and thus have broadly conserved roles in root-infecting fungi.

## Materials and Methods

### Fungal strains and transformants

Fungal strains used in this study are listed in table S2. All the generated knockouts are derivatives of *F. oxysporum* f. sp. *lycopersici* isolate 4287 (NRRL34936) or *F. oxysporum* Fo47 (NRRL54002). Strain cultures and storage were performed as described (11). Phenotypic analysis of colony growth was done as previously reported (56). Targeted gene replacement with the hygromycin resistance cassette and complementation of the mutants by co-transformation with the phleomycin resistance cassette were performed as previously reported (56). Oligonucleotides used to generate PCR fragments for knockout generation by gene replacement, and complementation of mutants are listed in table S3. *F. oxysporum* gene data are available at the Fungal and oomycete Informatics Resources (FungiDB) under the following accession numbers: *erc1*, FOXG_11583; *erc2*, FOXG_04534; *erc3*, FOXG_16902; *erc4*, FOXG_08211; *ppi*, FOXG_08379.

### Plant growth conditions and infection assays

Tomato seeds (*S. lycopersicum* cv. Monica, Syngenta) were surface sterilized in 1% sodium hypochlorite for 30 mins and potted in vermiculite (Projar, Barcelona, Spain). Seedlings were grown in a growth chamber maintained at following conditions (15/9 h light/dark cycle, 28 °C). *M. polymorpha* accession Takaragaike-1 (Tak-1; male) was used for generating *Marchantia* thalli. *M. polymorpha* gemmae were grown on plates of half Gamborg’s B5 medium as described before (57).

Tomato root infection assays with *F. oxysporum* were performed as previously described (58) using a dipping protocol with 5 × 10^6^ *F. oxysporum* microconidia ml^−1.^ Survival was recorded daily and mortality curves were plotted by the Kaplan–Meier method and compared among groups using the log-rank test. *M. polymorpha* infection with *F. oxysporum* was performed as described (47).

### *G. mellonella* pathogenicity assays

Infection assays in *G. mellonella* larvae were performed as described previously (51). Briefly, a Burkard Auto Applicator (Burkard Manufacturing, UK) with a 1ml syringe was used to inject 8 μl of the microconidial suspension (2×10^7^ microconidia ml^-1^ in 1x PBS) into the hemocoel. Injected larvae were incubated in ventilated glass bottles at 30 °C and survival was recorded daily. Larvae were considered dead when they displayed no movement and were melanised (fig. S7F). Mortality curves were plotted by the Kaplan–Meier method and compared among groups using the log-rank test. The experiment was performed three times with similar results. Data presented are from one representative experiment.

### Generation of Fol-mClover3 or Fo47-mClover3 or Foc-mClover3-tagged *F. oxysporum* transformants

Plasmid pUC57 backbone carrying three copies of a *F. oxysporum* codon-optimized mClover3 (59) gene (Fo*-*mClover3), followed by three copies of the FLAG octapeptide tag coding region (3x*FLAG*) and driven by the *Aspergillus nidulans gpdA* promoter and the *SV40* late polyadenylation signal was synthesized by ProteoGenix (Schiltigheim, France). Codon-optimization of mClover3 was performed in accordance with *F. oxysporum* f. sp. *lycopersici* codon usage and GC content data retrieved from the Codon Usage Database (http://www.kazusa.or.jp/codon/).

Fo-mClover3-labeled strains of *F. oxysporum* f. sp. *lycopersici* (NRRL 34936), *F. oxysporum* f. sp. *cubense* (NRRL54006) and the *F. oxysporum* biocontrol isolate Fo47 (NRRL54002) were obtained by co-transforming fungal protoplasts with a Fo-mClover3 expression cassette (amplified from pUC57-Fo-mClover3 plasmid with primers gpdA-15b + SV40 Rev) and a hygromycin resistance cassette amplified from plasmid pAN7-1 (60) with primers gpdA-15b + Trpc8B, as previously described (12). Cytoplasmic Fo-mClover3 expression was observed and quantified in at least twenty independent transformants using a Zeiss Axio Imager M2 microscope (Zeiss, Barcelona, Spain) equipped with an Evolve Photometrics EM512 digital camera (Photometrics Technology, Tucson, AZ, United States) and a GFP filter set (BP 450/490, FT 510, LP 515). Fungal transformants showing brightest fluorescence were used in subsequent microscopy analysis.

### Laser scanning confocal microscopy

Laser Scanning confocal microscopy was performed using a Zeiss 880 Confocal microscope with Airyscan. *S. lycopersicum* roots or *M. polymorpha* thalli inoculated with fluorescent transformants of wild-type Fol4287, Foc54006 (TR4), Fo47 or different knockout derivatives thereof expressing cytoplasmic Fo-mClover3 were visualized at an excitation of 488 nm and emission detected at 495-540 nm. To visualize plant cell walls, samples were co-stained by 15 min incubation in 2 mg ml^-1^ propidium iodide (PI) in water in the dark for 15 mins before imaging. PI fluorescence was visualized at an excitation of 561 nm, and emission detected at 570-640 nm.

### Sample preparation for TEM/ SEM analysis

Sample preparation for transmission electron microscopy (TEM) was carried out according to a protocol previously reported with slight modifications (61). Briefly, roots were fixed for 90 min with 2.5% glutaraldehyde in 0.06M Sorensen phosphate buffer at pH 7.2. After 4 washes of 10 min each in the same buffer, the samples were post-fixed in 1% osmium tetroxide for 90 min. Samples were then rinsed 4 times 10 min each in buffer and dehydrated in a graded series of increasing concentrations of acetone (50%, 70%, 90%, and 100%) for 20 min per concentration. After dehydration samples were gradually infiltrated with increasing concentrations of EMBed812 resin (30%, 60%, and 100%) mixed with acetone for a minimum of 3h per step. Finally, samples were embedded in pure, fresh EMBed812 resin and polymerized for 48 h at 60°C. Ultrathin sections (80 nm) were cut with a Leica EM Ultracut UC7 ultramicrotome (Leica Microsystems, Vienna, Austria), post-stained 5 minutes with 1% (w/v) lead citrate dissolved in 0.6 M NaOH and subsequently 15 min with 2% (w/v) uranyl-acetate dissolved in distilled water. Sections were analyzed using a JEM1010 transmission electron microscope (JEOL, Tokyo, Japan).

For SEM, sample preparation was carried out as described (62) with slight modifications. Small pieces (1mm^2^) were cut from *Marchantia* thalli and fixed for 90 min with 2.5% glutaraldehyde in 0.06M Sorensen phosphate buffer at pH 7.2. After 4 washes of 10 min each in the same buffer, the samples were dehydrated in a graded series of increasing concentrations of ethanol (50 %, 70 %, 90 %, and 100 %) for 20 min per concentration. Dehydrated samples were critical point dried (Leica EM CPD 300; Leica Microsystems) using a customized program for plant leaves of about 80 min duration (settings for CO2 inlet: speed=medium & delay=120s; settings for exchange: speed=5 & cycles=18; settings for gas release: heat=medium & speed=medium). Samples were then mounted on aluminum stubs with carbon tape and sputter coated with 10 nm iridium (Leica EM ACE 600, Leica Microsystems) and imaged using a FEI Versa 3D scanning electron microscope (FEI, Hillsboro, OR, USA) under high vacuum condition.

### Quantification of gene expression and fungal burden by real-time quantitative PCR (qRT-PCR)

Real time (RT) qPCR for quantification of gene expression or of fungal biomass in *S. lycopersicum* or *M. polymorpha* plants was performed as described previously (47). Briefly, RNA was extracted using the Tripure RNA isolation reageant (Roche, Spain) with the DNase treatment (Roche, Spain). Reverse transcription was carried out using the Transcriptor Universal cDNA Master (Roche, Spain). qRT-PCR was performed using CFX96 Touch Real-Time PCR (Bio-Rad). Cycling conditions were 10 mins at 95°C followed by 40 cycles of 10 s at 95°C, 10 s at 62°C, and 20 s at 72°C. Data were analyzed using the ΔΔCt method (63) by calculating the ratio of the plant housekeeping genes *SlGapdh* (tomato) or *MpEF1a* (*M. polymorpha*) (64) versus the Fol4287-specific *six1* gene (*FOXG_16418*) or the Fo47-specific gene on Chr7 (*FOBG_10856*)^42^ to calculate the fungal burden. Expression profiling of the ERCs was carried out by using Fol4287-peptidyl prolyl isomerase gene (*FOXG_08379)* (47). Moreover, the *S. lycopersicum* defense related genes were tested by using the primers from previous report (65). For expression profiling of *V. dahliae erc* genes VdGAPDH was used as a reference (66). All primers used for the RT-qPCR analysis are listed in table S3.

### Isolation of tomato apoplastic fluid (AF) and fungal culture filtrate

For isolation of AF, roots of 2-week old tomato plants roots were used at 72h after inoculation. AF isolation was carried out as previously described (67) with minor modification. Briefly, roots were thoroughly washed in running water and cut into pieces of 2cm length. Samples were vacuum infiltrated with deionized water 3 times for 15 min at 100 mbar with 5 min atmospheric pressure breaks. Roots were then thoroughly tap dried on a tissue paper. Bundled infiltrated roots were centrifuged in 10 ml syringe barrels at 2,200 r.p.m. for 15 min at 4 °C. Pooled AF was flash frozen and stored at −80 °C.

To process the protein samples for LC-MS, the isolated AF was concentrated on an Amicon Ultra 0.5ml -3KDa cutoff (Merck Millipore, Germany) and the concentrate was further resolved in 20μl of 1× SDS sample buffer and 25 μl were used. Proteins were analysed by SDS–polyacrylamide (10%) gel electrophoresis with a short run, followed by Coomassie blue staining. Protein bands of 1cm block on the resolving gel were eluted and tested to identify proteins in AF. An In-gel Trypsin digestion was carried out and samples were analysed by liquid chromatography-mass spectrometry (LC-MS). Identification of tomato and *F. oxysporum* proteins was carried out at the RTP, Proteomics Facility, University of Warwick, UK.

Filtrates from axenic cultures of *F. oxysporum* secreted proteins were obtained as previously described (53). Briefly, 3-day-old cultures of Fol4287 grown in PDB or in minimal medium (MM) supplemented either with crushed roots or sucrose as carbon source were harvested by filteration, first through a cheesecloth membrane and then through a 0.45μm syringe filter (Merck Millipore, Germany). Proteins were precipitated by adding 10% (v/v, final concentration) trichloroacetic acid, incubating at −20°C overnight and subsequent centrifugation at 20,000 *g*. Pelleted secreted proteins were resolved in 50 μl of 1× SDS sample buffer and 25 μl were used for SDS–PAGE analysis.

### Western Blot Analysis

To exclude cytoplasmic contamination a Western blot was performed on the isolated AF along with the total root extract as positive control with tubulin antibody. For Western blot, proteins from SDS-PAGE gels were transferred onto nitrocellulose membranes using the Transblot Turbo RTA Midi Transfer kit (BioRad, USA). Mouse anti-α-tubulin antibody (Sigma-Aldrich; #T9026) was used at a 1:5,000 dilution to determine leakage of cytoplasmic contamination into the apoplastic fluid. The membrane was visualized using the ECL Select Western blotting detection reagent (GE Healthcare, Chicago, IL, USA) in a LAS-3000 detection system (Fujifilm, Barcelona, Spain).

### Phylogenetic analysis and tree construction

Protein sequences of putative effectors were obtained from the Department of Energy-Joint Genome Institute MycoCosm database (https://mycocosm.jgi.doe.gov/mycocosm/home). A blastp search was performed using ERC1, ERC2 and ERC3 against the MycoCosm database. For phylogenetic tree construction, protein sequences were selected based on sequence homology. Multiple sequence alignment was performed using MUSCLE (v3.8.425) with default parameters over 100 iterations (68). The phylogenetic tree was constructed with RAxML (v8.2.12) using the PROTGAMMAWAG model for protein sequence alignment (69). Bootstrap support was determined for all the phylogenetic trees with a convergence test to confirm sufficient sampling.

### RNAseq and data analysis

Roots of 2-week-old Tomato seedlings were collected at 1, 2, 3 or 7 dpi with Fol4287. The axenic growth control was generated as follows; fresh microconidia of Fol4287 obtained from a 3-d-culture in PDB as described (56) were inoculated in 200 ml fresh PDB at a concentration of 2×10^6^ microconidia ml^−1^. After 13 h incubation at 28°C, the germlings were harvested by filteration through a cheesecloth membrane, resuspended in 200 ml MM containing NaNO_3_ as nitrogen source and incubated an additional 5 h at 28°C. Germlings were then harvested and flash frozen for RNA isolation. Total RNA was isolated using the RNeasy Plant Mini Kit (Qiagen, Germany) and treated with DNAse I usind the Turbo DNA Free Kit (Invitrogen, Germany) according to the manufacturer’s instructions. RNA sequencing was performed by Novogene, UK. For library preparation mRNA was captured through poly-A enrichment on the total RNA, and a TruSeq RNA Library Preparation Kit (Illumina, USA) was used to build the libraries according to the manufacturer’s protocol. Libraries were sequenced on a NovaSeq6000 sequencing platform (Illumina). Paired-end 150-bp reads were obtained for each RNAseq library.

### Read mapping and differential expression analysis

Transcript quantification was performed with Salmon (70). RNASeq paired-end read datasets were quasi mapped against the reference transcriptome of *F. oxysporum* (GCF_000149955.1_ASM14995v2_rna.fna, obtained from NCBI RefSeq). Adaptor-trimmed reads have been uploaded to the ArrayExpress database. Differential gene expression analysis on gene and transcript level was analyzed using DESeq2 (1.28.1) (71), following a pair-wise comparison between *F. oxysporum* samples *in planta* as compared to the axenic growth control. GO-terms for *F. oxysporum* were obtained by processing the GCF_000149955.1 proteins with annotF (https://github.com/gemygk/AnnotF), wrapping Blast (72), Blast2GO (73) and InterProScan (74) GO-term analysis was done in R using the topGO package (75) (2.40.0), by testing for enrichment in each infection timepoint as compared to the axenic growth control. Differentially expressed genes (DEGs) (absolute LFC [log2 fold change] > 2 and adjusted p value <0.01) were used to create volcano plots and hierarchical clustering of samples. Secreted proteins from the DEGs were predicted with SignalP5.0 (76), using a minimal signal peptide threshold of 0.75. Heatmaps for the differentially expressed secreted genes were generated with the R heatmap package (1.0.12) using variance-stabilized counts median-centered by gene. Scripts used to analyze RNA-seq datasets and visualize differentially expressed genes are available at https://github.com/cschu/redkar_et_al_erc. Expression plots for the effectors of interest and for the amino acid transporters enriched in GO processes were done in Python by plotting the log2 expression values of the candidate gene on the y-axis versus, time in dpi on the x-axis.

Chromosomal locations of DEGs were visualized using a custom Python script https://github.com/cschu/chrom_plot. To this aim, DEGs were assigned to three different groups: early expressed genes (average of all 3 expression values for 7 dpi/average of all 9 expression values for 1-3 dpi > 0.25) and represented in red; late expressed genes (average of all 9 expression values for 1-3 dpi/average of all 3 expression values for 7 dpi < 0.25) and represented in blue; or expressed at all timepoints (DE at 7 dpi and at least one additional timepoint) and represented in green.

### Targeted knockouts of the *F. oxysporum erc* genes

Targeted gene replacement with the hygromycin resistance cassette and complementation of the mutants by co-transformation with the phleomycin resistance cassette were performed as reported previously (56). Oligonucleotides used to generate polymerase chain reaction (PCR) fragments for gene replacement, complementation or identification of mutants are listed (table S3). PCR reactions were performed with a High Fidelity Phusion Polymerase (NEB, Germany), using an Mini personal thermal cycler (BioRad).

### Statistical analysis

No statistical methods were used to predetermine sample size. Statistical analysis was carried out using the GraphPad Prism 9.0 software (San Diego, USA) and the data were plotted using the same tool. For statistical analysis, all data were tested with a non-parametric or mixed one-way ANOVA analysis followed by Bonferroni’s multiple comparison test or Unpaired t-test for statistical significance. Data points with different letters indicate significant differences of *P <* 0.05 for Bonferroni’s test results. Data points are plotted onto the graphs, and the number of samples or the description of error bars are indicated in the corresponding figure legends. (\**P* ≤ 0.05; \*\**P* ≤ 0.01; \*\*\**P* ≤ 0.001; \*\*\*\**P* ≤ 0.0001; otherwise, not significant) between samples. For the Kaplan-Meier plots showing comparison between percent survival of plants, log-rank (Mantel-Cox) test was performed to calculate the statistical significance (p<0.05) of survival compared to wild type infected plants.

## Supporting information

Suppmentary information

## Acknowledgements

We thank M.I.G. Roncero (Universidad de Córdoba, Spain) for critical reading of the manuscript. We thank the Central Service for Research Support (SCAI) of the University of Córdoba for confocal microscopy facility. The high-performance computing resources and services in this work were supported by the Earlham Institute Scientific Computing group alongside the Norwich BioScience Institutes Partnership Computing infrastructure for Science (CiS) group.

## Funding

This work was supported from the Spanish Ministry of Science and Innovation (MICINN, grant PID2019-108045RB-I00) to A.D.P. A.R. and M.S. were supported by the European Union’s Horizon 2020 research and innovation program under the Marie Skłodowska-Curie grant agreements No. 750669 and 797256. A.R. also acknowledges funding from Juan de la Cierva Incorporación grant from the Spanish Research Agency (IJC2018-038468-I). C.S. was supported by BBSRC strategic funding, Core Capability Grant BB/CCG1720/1, BBS/E/T/000PR9816. Research in R.S. lab was funded by the Spanish Ministry for Science and Innovation grant PID2019-107012RB-100 (MICINN/FEDER).

## Author Contributions

A.R. and A.D.P. conceptualized the work, designed the experiments and supervised the conducted research. A.R., M.S., B.Z., Y.K.G., M.S.L.B., S.G.I., G.V. and D.T. carried out the experiments and analysed the data. C.S. performed all the bioinformatic analysis. R.S. and S.G.I contributed in sample material and gave intellectual input for the *Marchantia* related experiments. A.R. and A.D.P. wrote the manuscript. All authors participated in reviewing and editing the final version of the manuscript.

## Competing interests

The authors declare that they have no competing interests.

## Data and materials Availability

All data needed to evaluate the conclusions in the paper are present in the paper and/or the Supplementary Materials. The mass spectrometry proteomics data have been deposited to the ProteomeXchange Consortium via the PRIDE partner repository. The RNA-seq datasets generated and analysed in the current study have been deposited in the ArrayExpress database at EMBL-EBI (www.ebi.ac.uk/arrayexpress) (Accession numbers will be available upon acceptance of publication).

## Notes

### Competing Interest Statement

The authors have declared no competing interest.

