## Supplementary material for "Conserved secreted effectors determine endophytic growth and multi-host plant compatibility in a vascular wilt fungus": Suppmentary information

**This PDF file includes:**

Figs. S1 to S7
Tables S1 to S4
References (S1 to S3)

**Other Supplementary Materials for this manuscript include the following:**
Data S1 to S10

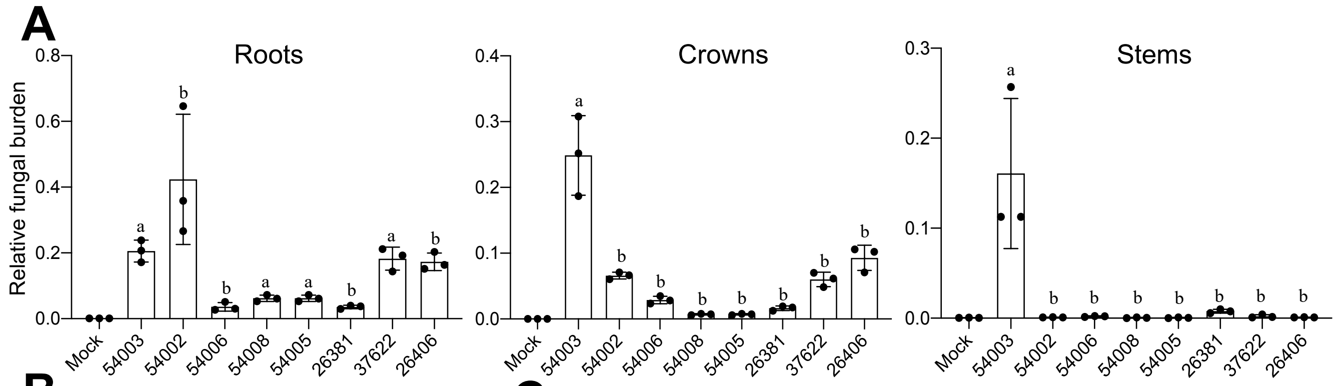

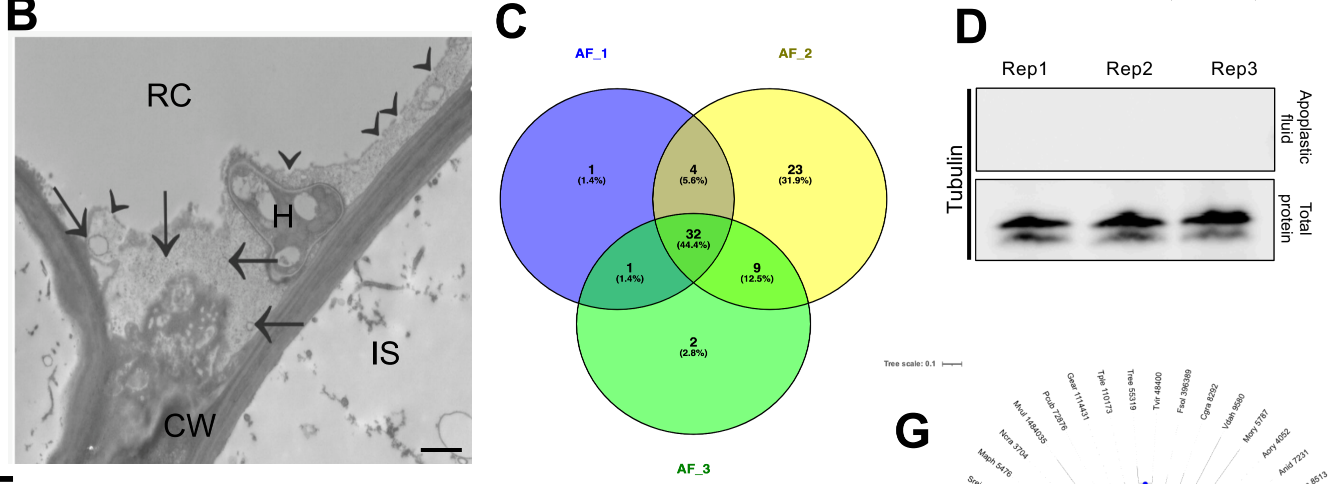

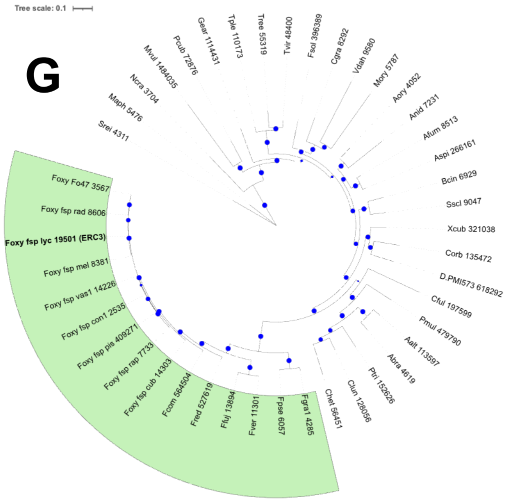

**E**

| Gene Number | Abbreviation | Mol. weight | Signal Peptide | No of cysteines | Predicted domains |
|---|---|---|---|---|---|
| FOXG_11583 | ERC1 | 23 kDa | Yes | 6 | Expansion like superfamily; cellulose binding protein |
| FOXG_04534 | ERC2 | 28 kDa | Yes | 15 | lytic polysaccharide monooxygenase (LPMO) Domain |
| FOXG_16902 | ERC3 | 51 kDa | Yes | 8 | Alpha L arabinofuranosidase |
| FOXG_08211 | ERC4 | 34 kDa | Yes | 3 | LPMO and Glycosyl hydrolase family 61 |

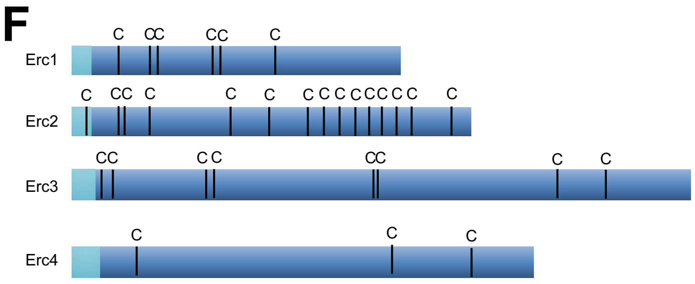

**Fig. S1. Fo colonizes roots of host and non-host plants and secretes conserved core effectors.** (**A**) Fungal burden in roots, crowns and stems of tomato plants was measured by real time (qRT-qPCR) of the Fo *actin* gene using total genomic DNA extracted at 35 days after dip-inoculation with the indicated Fo isolates belonging to different ff. spp. (listed in table S1). Fo f.sp. *lycopersici* race 3 (54003) was used as a reference for DNA quantification of host to non-host interaction to all other ff.sp. tested. $n_{i.ex} = 3$; 10 plants/treatment. Data are presented as mean values ± s.d. from three independent experiments. Different letters indicate statistically significant differences according to one-way ANOVA followed by Bonferroni´s multiple comparison test ($P < 0.05$). (**B**) TEM micrograph showing a hypha of Fol4287 (H) within a tomato root cell (RC). Note that the infected plant cell shows loss of plasma membrane integrity (arrowheads) and an accumulation of vesicles (arrows). CW; plant cell wall, IS; intercellular space. Scale bar, 1 µm. (**C**) Venn diagram showing Fol4287 proteins identified in the three biological replicates of tomato root apoplastic fluid (AF). Among a total of 72 proteins, 32 were detected in all replicates. (**D**) Western blot with anti-tubulin antibody performed with AF or crushed root total protein extracts (positive control) to ensure the absence of leakage of intracellular Fo proteins into AF samples used for mass spectrometry analysis. The three lanes correspond to the different replicates tested. (**E**) Table summarizing the characteristics of the four identified ERC effectors. (**F**) Schematic representation of the four identified ERC effector proteins. Light and dark blue boxes indicate predicted signal peptide and mature protein, respectively. The relative positions of cysteine residues (C) are indicated. (**G**) Maximum likelihood phylogenetic tree based on the aligned amino acid sequences of the *FOXG_16902* (ERC3) protein. Size of blue dots represents bootstrap support for the branch with maximum 100 bootstraps. Fungal species included in the analysis are listed in table S4.

adj.pval: <0.05 (Significant) — 0.05 — >0.05 (Non-significant)

# A

#### 1dpi

| | Target | Log2FC | adj.pval |
|---|---|---|---|
| FoxG_14489 | Exo beta 1,3, Galactanase | 11,994 | 1,29E-14 |
| FoxG_15793 | hypothetical protein | 10,513 | 1,24E-10 |
| FoxG_05669 | uncharacterized protein | 10,309 | 9,03E-10 |
| FoxG_13032 | choline dehydrogenase | 10,043 | 1,59E-44 |
| FoxG_13249 | pectate lyase | 9,204 | 1,62E-06 |
| FoxG_17428 | Beta xylanase | 8,531 | 3,11E-100 |
| FoxG_07464 | Copper amineoxidase | 7,664 | 6,55E-239 |
| FoxG_16880 | Endoglucanase | 7,601 | 0,000156 |
| FoxG_16902 | Alpha L arabinofuranosidase | 7,565 | 2,14E-06 |
| FoxG_11739 | Pectate lyase | 7,400 | 5,50E-13 |
| FoxG_13638 | carbonic anhydrase | 7,234 | 5,11E-59 |
| FoxG_09125 | Beta glucosidase | 6,813 | 8,17E-19 |
| FoxG_08316 | uncharacterized protein | 6,679 | 8,75E-53 |
| FoxG_11736 | uncharacterized protein | 6,612 | 8,83E-38 |
| FoxG_11583 | Uncharacterized protein | 6,555 | 7,12E-130 |
| FoxG_17608 | Alpha Galactosidase | 6,087 | 0,003906 |
| FoxG_00928 | Aldehyde dehydrogenase | 6,055 | 5,03E-69 |
| FoxG_13188 | Xyloglucanase | 5,765 | 1,26E-05 |
| FoxG_17404 | Murein transglycosylase | 5,746 | 0,013882 |
| FoxG_09643 | Glucanase | 5,746 | 0,010377 |
| FoxG_01352 | uncharacterized protein | 5,746 | 0,026382 |
| FoxG_13239 | Alpha galactosidase | 5,305 | 0,013732 |
| FoxG_14695 | Endopolygalacturonase | 4,902 | 0,011246 |
| FoxG_04966 | hypothetical protein | 4,881 | 1,54E-33 |
| FoxG_08211 | hypothetical protein | 4,557 | 1,44E-51 |
| FoxG_10463 | uncharacterized protein | 4,394 | 0,041291 |
| FoxG_04408 | nucleotidase | 4,160 | 5,36E-41 |
| FoxG_09029 | uncharacterized protein | 4,044 | 1,04E-62 |
| FoxG_14528 | oxidoreductases | 3,856 | NA |
| FoxG_09688 | uncharacterized protein | 2,882 | 0,151288 |
| FoxG_04534 | uncharacterized protein | 1,453 | 0,694003 |
| FoxG_08027 | Glucoamylase | 1,334 | 0,001356 |
| FoxG_00480 | Glucanase | 1,231 | 0,112049 |
| FoxG_12261 | uncharacterized protein | 1,220 | 0,328145 |
| FoxG_02142 | Beta glucosidase | 0,868 | 0,205224 |
| FoxG_13353 | Alpha L fucosidase | 0,720 | 0,758371 |
| FoxG_17211 | uncharacterized protein | 0,705 | NA |
| FoxG_15833 | hypothetical protein | 0,424 | 0,610861 |
| FoxG_11102 | Aminopeptidase | -0,962 | 0,009903 |
| FoxG_03276 | uncharacterized protein | -1,318 | 1,05E-06 |
| FoxG_14546 | hypothetical protein | -1,429 | 0,350978 |
| FoxG_11203 | Glucose 6 phosphate epimerase | -1,552 | 0,016363 |
| FoxG_09264 | Trehalase | -1,894 | 0,000434 |
| FoxG_15430 | peroxidase | -2,462 | 4,00E-22 |
| FoxG_02233 | Triosephosphate isomerase | -3,441 | 1,33E-33 |
| FoxG_13120 | uncharacterized protein | -3,687 | 0,200919 |

$Log_2FC$: 11.99 (Up-regulated) — 0 — -3.69 (Down-regulated)

#### 2dpi

| | Target | Log2FC | adj.pval |
|---|---|---|---|
| FoxG_16880 | Endoglucanase | 14,587 | 4,11E-21 |
| FoxG_09643 | Glucanase | 14,304 | 1,00E-19 |
| FoxG_17404 | Murein transglycosylase | 13,445 | 9,77E-16 |
| FoxG_05669 | uncharacterized protein | 13,386 | 5,31E-17 |
| FoxG_14489 | Exo beta 1,3, Galactanase | 13,075 | 1,42E-17 |
| FoxG_13249 | pectate lyase | 12,804 | 2,82E-13 |
| FoxG_10463 | uncharacterized protein | 12,387 | 8,80E-19 |
| FoxG_13188 | Xyloglucanase | 11,772 | 2,79E-32 |
| FoxG_01352 | uncharacterized protein | 11,303 | 7,52E-09 |
| FoxG_09688 | uncharacterized protein | 11,077 | 9,51E-24 |
| FoxG_15793 | hypothetical protein | 10,930 | 7,62E-12 |
| FoxG_16902 | Alpha L arabinofuranosidase | 10,839 | 6,30E-15 |
| FoxG_11739 | Pectate lyase | 10,708 | 3,98E-28 |
| FoxG_13032 | choline dehydrogenase | 10,595 | 2,73E-50 |
| FoxG_13239 | Alpha galactosidase | 9,224 | 1,28E-09 |
| FoxG_17428 | Beta xylanase | 9,130 | 1,41E-118 |
| FoxG_11736 | uncharacterized protein | 8,948 | 1,65E-78 |
| FoxG_13638 | carbonic anhydrase | 8,680 | 1,87E-94 |
| FoxG_14695 | Endopolygalacturonase | 7,680 | 5,11E-06 |
| FoxG_09125 | Beta glucosidase | 7,327 | 7,88E-23 |
| FoxG_00480 | Glucanase | 7,304 | 1,09E-50 |
| FoxG_07464 | Copper amineoxidase | 6,976 | 4,64E-197 |
| FoxG_11583 | Uncharacterized protein | 6,713 | 2,14E-137 |
| FoxG_08316 | uncharacterized protein | 5,951 | 5,12E-42 |
| FoxG_17608 | Alpha Galactosidase | 5,868 | 0,003730 |
| FoxG_00928 | Aldehyde dehydrogenase | 5,762 | 1,21E-62 |
| FoxG_04966 | hypothetical protein | 5,035 | 5,29E-37 |
| FoxG_04408 | nucleotidase | 4,433 | 1,81E-50 |
| FoxG_04534 | uncharacterized protein | 4,245 | 0,125950 |
| FoxG_08211 | hypothetical protein | 4,169 | 1,78E-43 |
| FoxG_12261 | uncharacterized protein | 3,706 | 1,87E-05 |
| FoxG_08027 | Glucoamylase | 3,499 | 3,27E-23 |
| FoxG_09029 | uncharacterized protein | 3,377 | 7,28E-44 |
| FoxG_02142 | Beta glucosidase | 3,311 | 1,96E-13 |
| FoxG_14528 | oxidoreductases | 3,251 | NA |
| FoxG_13353 | Alpha L fucosidase | 3,148 | 0,020506 |
| FoxG_15833 | hypothetical protein | 0,306 | 0,704436 |
| FoxG_17211 | uncharacterized protein | 0,101 | NA |
| FoxG_14546 | hypothetical protein | -0,890 | 0,512613 |
| FoxG_11203 | Glucose 6 phosphate epimerase | -1,409 | 0,023054 |
| FoxG_15430 | peroxidase | -1,501 | 8,30E-10 |
| FoxG_09264 | Trehalase | -1,786 | 0,000139 |
| FoxG_11102 | Aminopeptidase | -1,859 | 1,07E-07 |
| FoxG_03276 | uncharacterized protein | -2,052 | 5,20E-15 |
| FoxG_13120 | uncharacterized protein | -2,706 | 0,342504 |
| FoxG_02233 | Triosephosphate isomerase | -3,781 | 2,12E-42 |

$Log_2FC$: 14.59 (Up-regulated) — 0 — -3.78 (Down-regulated)

#### 3dpi

| | Target | Log2FC | adj.pval |
|---|---|---|---|
| FoxG_16880 | Endoglucanase | 15,506 | 9,21E-24 |
| FoxG_09643 | Glucanase | 14,054 | 5,00E-19 |
| FoxG_17404 | Murein transglycosylase | 12,775 | 3,18E-14 |
| FoxG_15793 | hypothetical protein | 12,553 | 1,29E-15 |
| FoxG_14489 | Exo beta 1,3, Galactanase | 12,421 | 7,61E-16 |
| FoxG_10463 | uncharacterized protein | 12,196 | 3,48E-18 |
| FoxG_05669 | uncharacterized protein | 11,942 | 1,49E-13 |
| FoxG_13188 | Xyloglucanase | 11,469 | 1,27E-30 |
| FoxG_16902 | Alpha L arabinofuranosidase | 10,879 | 5,11E-15 |
| FoxG_13032 | choline dehydrogenase | 10,696 | 2,56E-51 |
| FoxG_13249 | pectate lyase | 9,951 | 4,97E-08 |
| FoxG_09125 | Beta glucosidase | 9,752 | 1,58E-41 |
| FoxG_01352 | uncharacterized protein | 8,744 | 2,83E-05 |
| FoxG_09688 | uncharacterized protein | 8,742 | 1,13E-14 |
| FoxG_11736 | uncharacterized protein | 8,489 | 2,89E-70 |
| FoxG_13239 | Alpha galactosidase | 8,348 | 8,87E-08 |
| FoxG_17608 | Alpha Galactosidase | 7,847 | 1,72E-05 |
| FoxG_00480 | Glucanase | 7,552 | 3,12E-54 |
| FoxG_13638 | carbonic anhydrase | 7,306 | 1,10E-63 |
| FoxG_11739 | Pectate lyase | 7,113 | 2,93E-12 |
| FoxG_17428 | Beta xylanase | 7,005 | 9,48E-65 |
| FoxG_07464 | Copper amineoxidase | 6,964 | 5,83E-197 |
| FoxG_08316 | uncharacterized protein | 6,443 | 2,27E-49 |
| FoxG_00928 | Aldehyde dehydrogenase | 6,284 | 1,77E-74 |
| FoxG_14695 | Endopolygalacturonase | 5,898 | 0,000879 |
| FoxG_11583 | Uncharacterized protein | 5,388 | 5,19E-87 |
| FoxG_04966 | hypothetical protein | 4,458 | 1,73E-28 |
| FoxG_02142 | Beta glucosidase | 4,441 | 1,73E-24 |
| FoxG_09029 | uncharacterized protein | 4,400 | 1,20E-75 |
| FoxG_08027 | Glucoamylase | 4,349 | 5,75E-36 |
| FoxG_14528 | oxidoreductases | 4,159 | NA |
| FoxG_04408 | nucleotidase | 3,346 | 2,91E-26 |
| FoxG_08211 | hypothetical protein | 3,316 | 3,20E-27 |
| FoxG_04534 | uncharacterized protein | 3,232 | 0,290040 |
| FoxG_13353 | Alpha L fucosidase | 1,928 | 0,292737 |
| FoxG_12261 | uncharacterized protein | 1,534 | 0,154427 |
| FoxG_15833 | hypothetical protein | 1,065 | 0,112328 |
| FoxG_09264 | Trehalase | 0,762 | 0,071560 |
| FoxG_14546 | hypothetical protein | 0,289 | 0,843161 |
| FoxG_17211 | uncharacterized protein | 0,048 | NA |
| FoxG_11102 | Aminopeptidase | -0,571 | 0,127354 |
| FoxG_11203 | Glucose 6 phosphate epimerase | -0,915 | 0,167722 |
| FoxG_13120 | uncharacterized protein | -1,124 | 0,704520 |
| FoxG_03276 | uncharacterized protein | -1,467 | 1,49E-08 |
| FoxG_15430 | peroxidase | -2,118 | 3,98E-18 |
| FoxG_02233 | Triosephosphate isomerase | -2,841 | 1,72E-27 |

$Log_2FC$: 15.51 (Up-regulated) — 0 — -2.84 (Down-regulated)

#### 7dpi

| | Target | Log2FC | adj.pval |
|---|---|---|---|
| FoxG_16880 | Endoglucanase | 14,935 | 3,83E-22 |
| FoxG_09643 | Glucanase | 14,776 | 4,87E-21 |
| FoxG_14489 | Exo beta 1,3, Galactanase | 13,958 | 4,99E-20 |
| FoxG_17404 | Murein transglycosylase | 12,466 | 1,15E-13 |
| FoxG_05669 | uncharacterized protein | 12,354 | 1,28E-14 |
| FoxG_13249 | pectate lyase | 12,351 | 1,91E-12 |
| FoxG_10463 | uncharacterized protein | 11,906 | 1,88E-17 |
| FoxG_15793 | hypothetical protein | 11,534 | 2,30E-13 |
| FoxG_13188 | Xyloglucanase | 11,356 | 3,72E-30 |
| FoxG_01352 | uncharacterized protein | 11,022 | 1,66E-08 |
| FoxG_16902 | Alpha L arabinofuranosidase | 10,497 | 3,65E-14 |
| FoxG_09688 | uncharacterized protein | 10,205 | 2,47E-20 |
| FoxG_11739 | Pectate lyase | 9,599 | 9,13E-23 |
| FoxG_11736 | uncharacterized protein | 9,481 | 1,80E-89 |
| FoxG_13239 | Alpha galactosidase | 9,370 | 3,93E-10 |
| FoxG_07464 | Copper amineoxidase | 8,535 | 0 |
| FoxG_13032 | choline dehydrogenase | 8,514 | 2,06E-32 |
| FoxG_09125 | Beta glucosidase | 8,479 | 1,51E-31 |
| FoxG_00480 | Glucanase | 7,942 | 3,91E-60 |
| FoxG_13638 | carbonic anhydrase | 7,835 | 7,39E-78 |
| FoxG_14695 | Endopolygalacturonase | 7,816 | 3,12E-06 |
| FoxG_17428 | Beta xylanase | 7,420 | 8,05E-78 |
| FoxG_13353 | Alpha L fucosidase | 6,357 | 5,47E-10 |
| FoxG_08316 | uncharacterized protein | 6,305 | 1,57E-47 |
| FoxG_17608 | Alpha Galactosidase | 6,288 | 0,000814 |
| FoxG_00928 | Aldehyde dehydrogenase | 6,060 | 2,58E-69 |
| FoxG_11583 | Uncharacterized protein | 5,958 | 1,21E-108 |
| FoxG_04966 | hypothetical protein | 5,728 | 7,71E-50 |
| FoxG_09029 | uncharacterized protein | 3,726 | 1,00E-54 |
| FoxG_14528 | oxidoreductases | 3,725 | NA |
| FoxG_02142 | Beta glucosidase | 3,395 | 1,02E-14 |
| FoxG_04408 | nucleotidase | 2,917 | 6,66E-22 |
| FoxG_08211 | hypothetical protein | 2,785 | 8,97E-20 |
| FoxG_12261 | uncharacterized protein | 2,549 | 0,004837 |
| FoxG_15833 | hypothetical protein | 1,625 | 0,006883 |
| FoxG_04534 | uncharacterized protein | 0,539 | 0,891019 |
| FoxG_09264 | Trehalase | 0,459 | 0,299811 |
| FoxG_08027 | Glucoamylase | -0,168 | 0,758552 |
| FoxG_14546 | hypothetical protein | -0,381 | 0,783059 |
| FoxG_11203 | Glucose 6 phosphate epimerase | -0,858 | 0,188487 |
| FoxG_03276 | uncharacterized protein | -0,974 | 0,000145 |
| FoxG_13120 | uncharacterized protein | -0,980 | 0,734549 |
| FoxG_11102 | Aminopeptidase | -1,012 | 0,002107 |
| FoxG_17211 | uncharacterized protein | -1,170 | NA |
| FoxG_15430 | peroxidase | -2,227 | 4,61E-21 |
| FoxG_02233 | Triosephosphate isomerase | -2,464 | 6,43E-23 |

$Log_2FC$: 14.94 (Up-regulated) — 0 — -2.46 (Down-regulated)

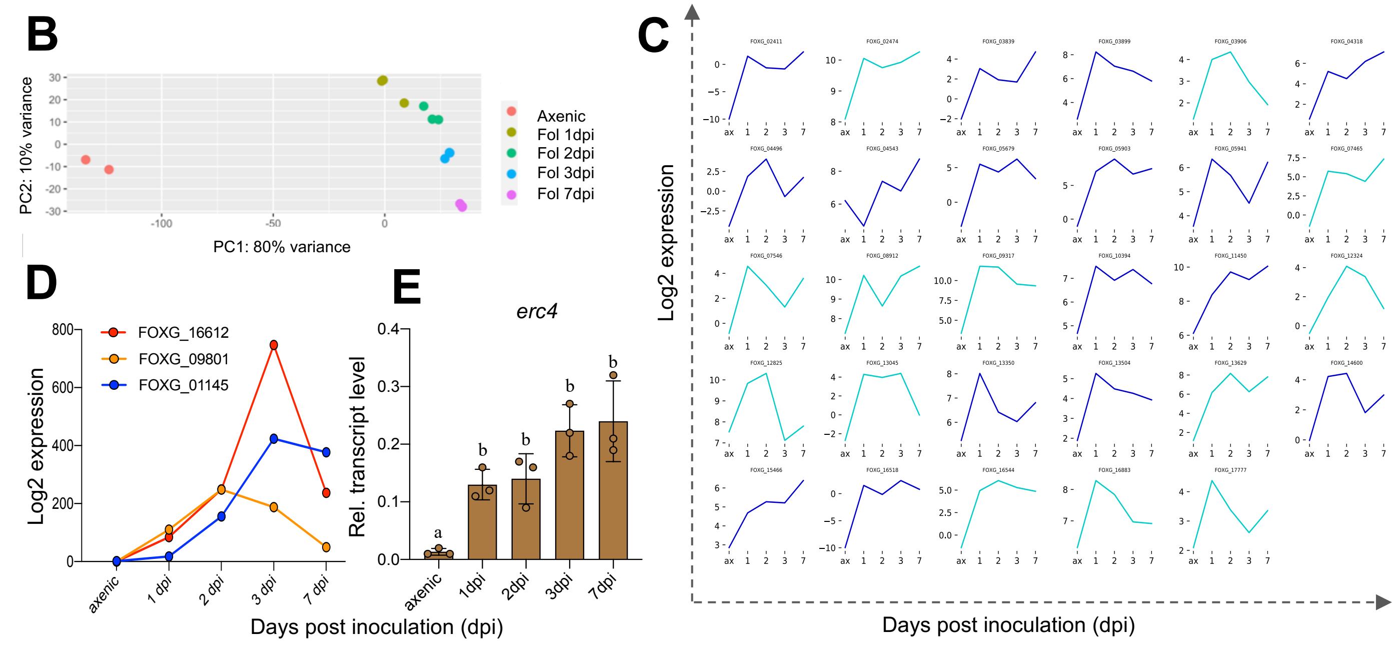

**Fig. S2. Transcriptional dynamics in Fo during early stages of root infection.** (**A**) Log2-transformed fold changes in transcript levels of genes encoding Fol4287 proteins detected in tomato root AF by MS analysis. Transcript levels of each gene were normalized against that detected in axenic culture. (adj. pval), adjusted *P* value. Benjamini–Hochberg FDR two-sided adjusted *P* value. $P < 0.05$ is considered significant. Red gradient, genes significantly upregulated in roots (positive log2-transformed fold change, $P < 0.05$). Blue gradient, genes significantly downregulated in roots. Green gradient, level of significance of the adjusted *P* value. The full list of differentially expressed genes is shown in Supplementary Datasets S6-S9. (**B**) Principal Component Analysis (PCA) plot showing transcriptional reprogramming of gene expression between axenic culture, the early (1,2,3 dpi) and the late (7 dpi) stages of infection. Red, Axenic; Olive, 1dpi; green, 2dpi; blue, 3dpi; pink, 7dpi. (**C**) Expression profiles (log2-transformed fold changes) of genes encoding amino acid transporters either lacking (blue) or containing a transmembrane domain (cyan), showing transcriptional upregulation during the early biotrophic infection phase. (**D**) Log2 expression counts showing upregulation of three genes encoding previously reported Fol4287 effectors belonging to metallo- (*FOXG_16612*) and serine proteases (*FOXG_09801, FOXG_01145*) during early stages of tomato root infection. (**E**) Relative transcript levels of gene *FOXG_08211* (*erc4*), were measured by qRT-PCR of cDNA obtained from Fol4287 grown in minimal medium (axenic) or from roots of tomato plants inoculated with Fol4287 at 1, 2, 3 or 7 dpi. Transcript levels were calculated using the threshold cycle (ΔΔCt) method and normalized to the Fol4287 peptidyl prolyl isomerase (*ppi*) gene. Error bars indicate standard deviation (s.d.); n = 3 biological replicates. Different letters indicate statistically significant differences according to one way ANOVA, Bonferroni´s multiple comparison test ($p < 0.05$).

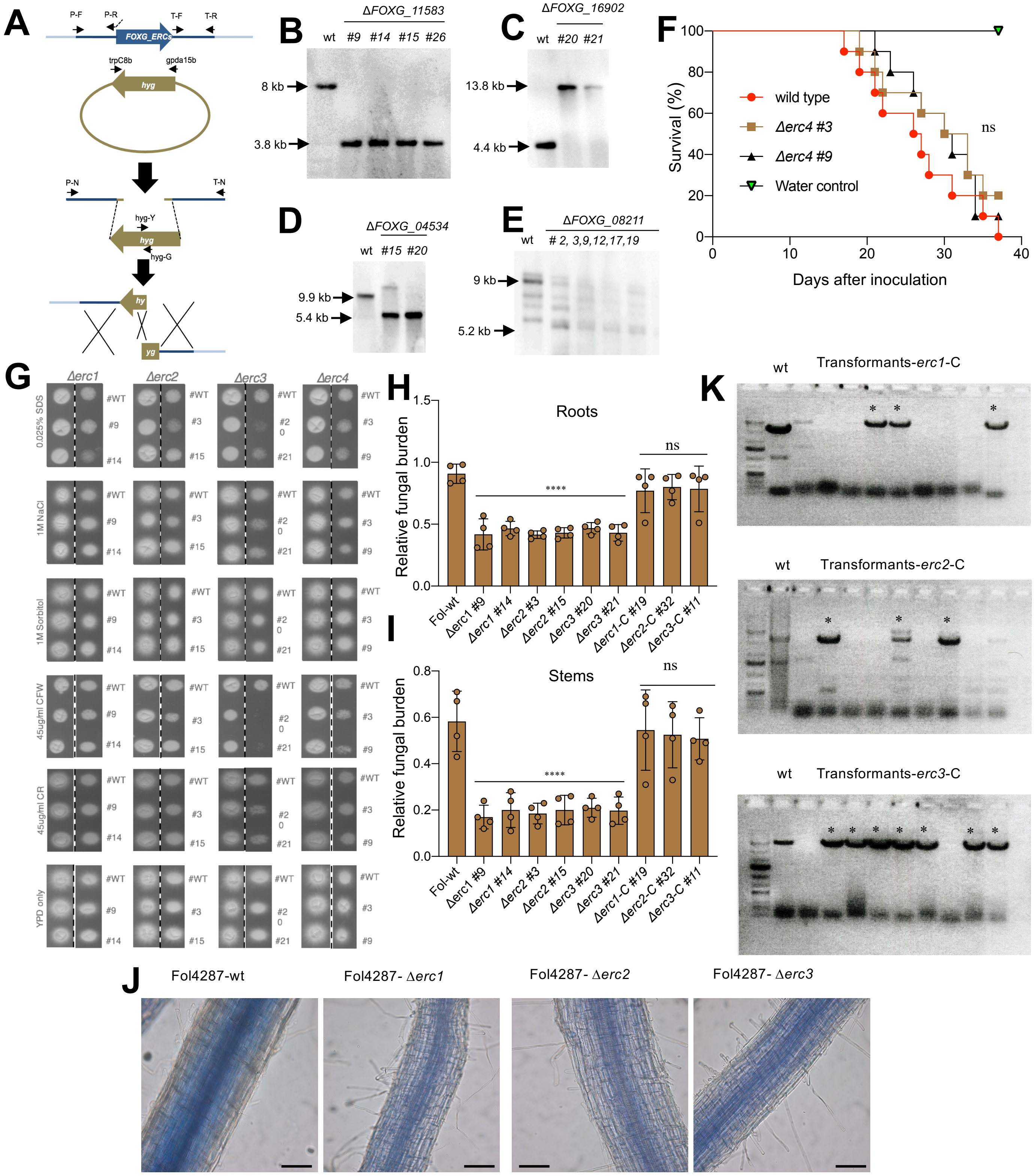
A
P-F
P-R
T-F
T-R
FOXG_ERCs
trpC8b
gpda15b
hyg
P-N
T-N
hyg-Y
hyg-G
hy
yg
B
ΔFOXG_11583
wt #9 #14 #15 #26
8 kb
3.8 kb
C
ΔFOXG_16902
wt #20 #21
13.8 kb
4.4 kb
D
ΔFOXG_04534
wt #15 #20
9.9 kb
5.4 kb
E
ΔFOXG_08211
wt
# 2, 3,9,12,17,19
9 kb
5.2 kb
F
Survival (%)
wild type
Δerc4 #3
Δerc4 #9
Water control
ns
Days after inoculation
G
Δerc1
Δerc2
Δerc3
Δerc4
0.025% SDS
1M NaCl
1M Sorbitol
45ug/ml CFW
45ug/ml CR
YPD only
#WT
#9
#14
#3
#15
#20
#21
H
Roots
Relative fungal burden
****
ns
Fol-wt
Δerc1 #9
Δerc1 #14
Δerc2 #3
Δerc2 #15
Δerc3 #20
Δerc3 #21
Δerc1-C #19
Δerc2-C #32
Δerc3-C #11
I
Stems
Relative fungal burden
****
ns
J
Fol4287-wt
Fol4287- Δerc1
Fol4287- Δerc2
Fol4287- Δerc3
K
wt
Transformants-erc1-C
Transformants-erc2-C
Transformants-erc3-C

**Fig. S3. Generation and phenotyping of ERC knockout mutants in Fo.** (**A**) Schematic diagram showing targeted replacement of *erc* genes using the split-marker technique. Gene replacement constructs were obtained by fusion PCR. Relative positions of the PCR primers used are indicated. hyg, hygromycin resistance casette. (**B** to **E**)**,** Southern blot analysis of genomic DNA of the wild type (wt) strain and independent *erc* knockout mutants, treated with restriction enzymes, separated on 0.7% agarose gels, transferred to a nylon membranes and hybridized with a DIG labelled DNA probes from the indicated genes. Molecular weights of the hybridizing bands are indicated on the left. (**F**) Kaplan-Meier plot showing the survival of tomato plants inoculated with the Fol4287 wild type strain or the indicated Δ*erc*4 mutants. Number of independent experiments = 3; 10 plants/treatment. Data shown are from one representative experiment. ns = non-significant versus Fol4287 according to log-rank test. (**G**) Stress response phenotypes of the different Fo Δ*erc* mutants. Drops containing 5 µl ($10^5$ and $10^4$) microconidia of the indicated strains were spotted on YDP medium alone or supplemented with SDS (0.025%), NaCl (1M), Sorbitol (1M), Calcofluor White (45 µg/mL) or Congo Red (45 µg/mL). Plates were incubated for two days at 28ºC. Images are representative of three independent replicates. (**H** and **I**) Fungal burden in tomato plants inoculated with the indicated Fo strains was measured by real time (qRT-PCR) of the Fo *actin* gene using total DNA extracted from roots (**H**) or stems (**I**) at 12 dpi. Fo DNA was calculated using the threshold cycle (ΔΔCt) method, normalized to the tomato *gadph* gene and expressed relative to Fol4287 in roots. Error bars indicate standard deviation (s.d.); n = 3 biological replicates. Asterisks indicate statistical significance versus Fol4287 (one way ANOVA, Bonferroni´s multiple comparison test, $p<0.05$). Experiments were performed three times with similar results. (**J**) Trypan blue staining of tomato roots inoculated with the Fol4287 wt and the indicated Δ*erc* deletion mutants. Scale bar = 0.1 mm. (**K**) Identification of complemented transformants of the indicated Δ*erc* deletion mutants, using PCR on genomic DNA of the indicated strains with primer pairs P-N and T-N indicated in (**A**). Presence of an amplification product indicates successful complementation of the knockout strain.

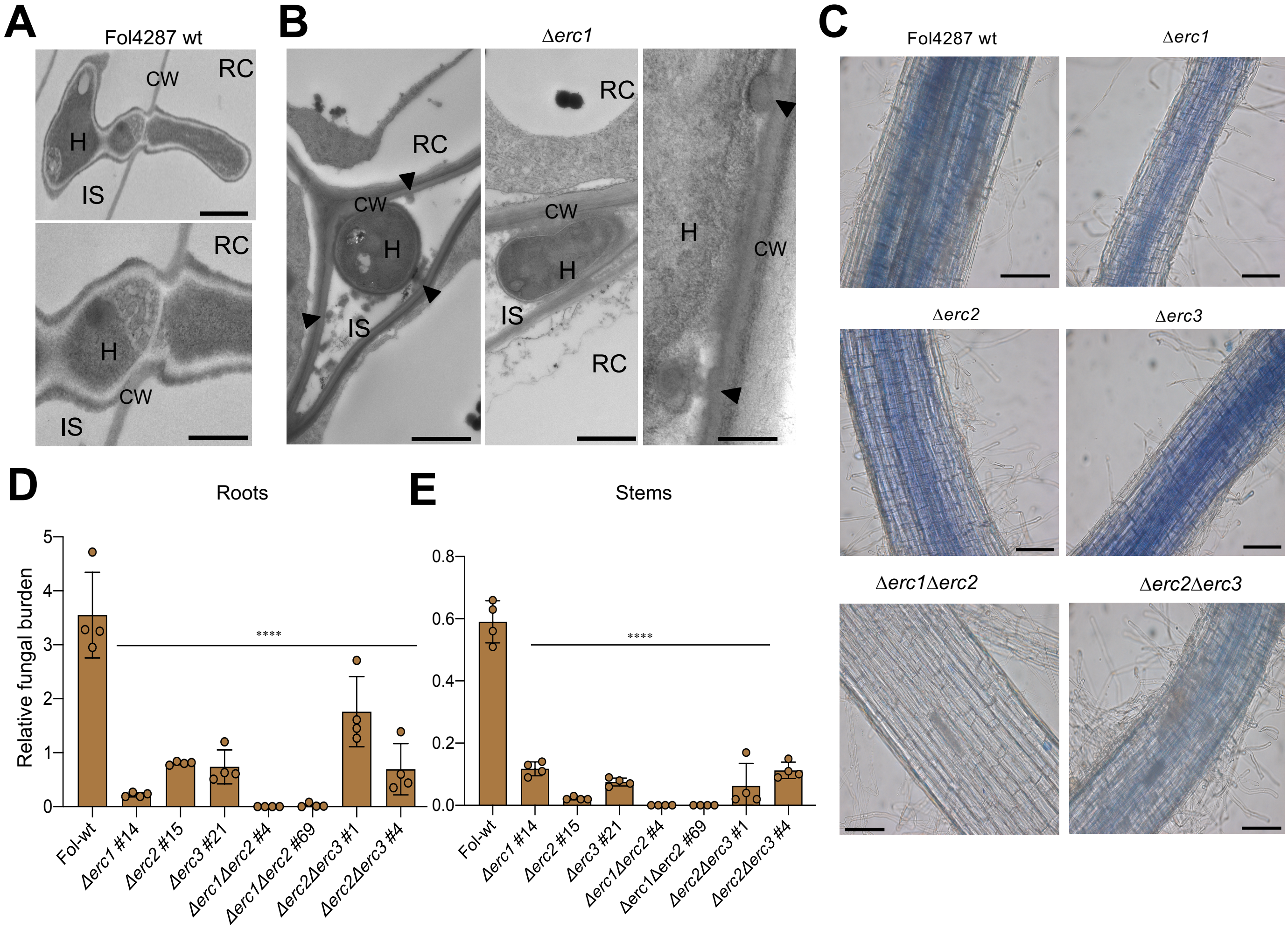

**Fig. S4. ERCs have a role in root colonization and virulence on the host plant.** (**A**) TEM micrographs showing hyphae (H) of Fol4287 growing between and penetrating into tomato root cells (RC). CW; plant cell wall, IS; intercellular space. Scale bars, 1 µm (top) and 0.5 µm (bottom). (**B**) TEM micrographs showing hyphae (H) of the Δ*erc1* mutant growing between tomato root cells (RC). Note that hyphae are encapsulated by an amorphous granular material and by protrusions of the plant cell wall (arrowheads). Scale bars, 0.5 µm. (**C**) Trypan blue staining of tomato roots inoculated with the Fol4287 wt and the indicated Δ*erc* deletion mutants. Scale bar = 0.1 mm. (**D** and **E**) Fungal burden in tomato plants inoculated with the indicated Fo isolates was measured by real time (qRT-PCR) of the Fo *actin* gene, using total DNA extracted from roots or stems at 12 dpi. Fo DNA was calculated using the threshold cycle (ΔΔCt) method, normalized to the tomato *gadph* gene. Error bars indicate standard deviation (s.d.); n = 3 biological replicates. Asterisks indicate statistical significance versus Fol4287 (one way ANOVA, Bonferroni´s multiple comparison test, $p<0.05$). Experiments were performed three times with similar results.

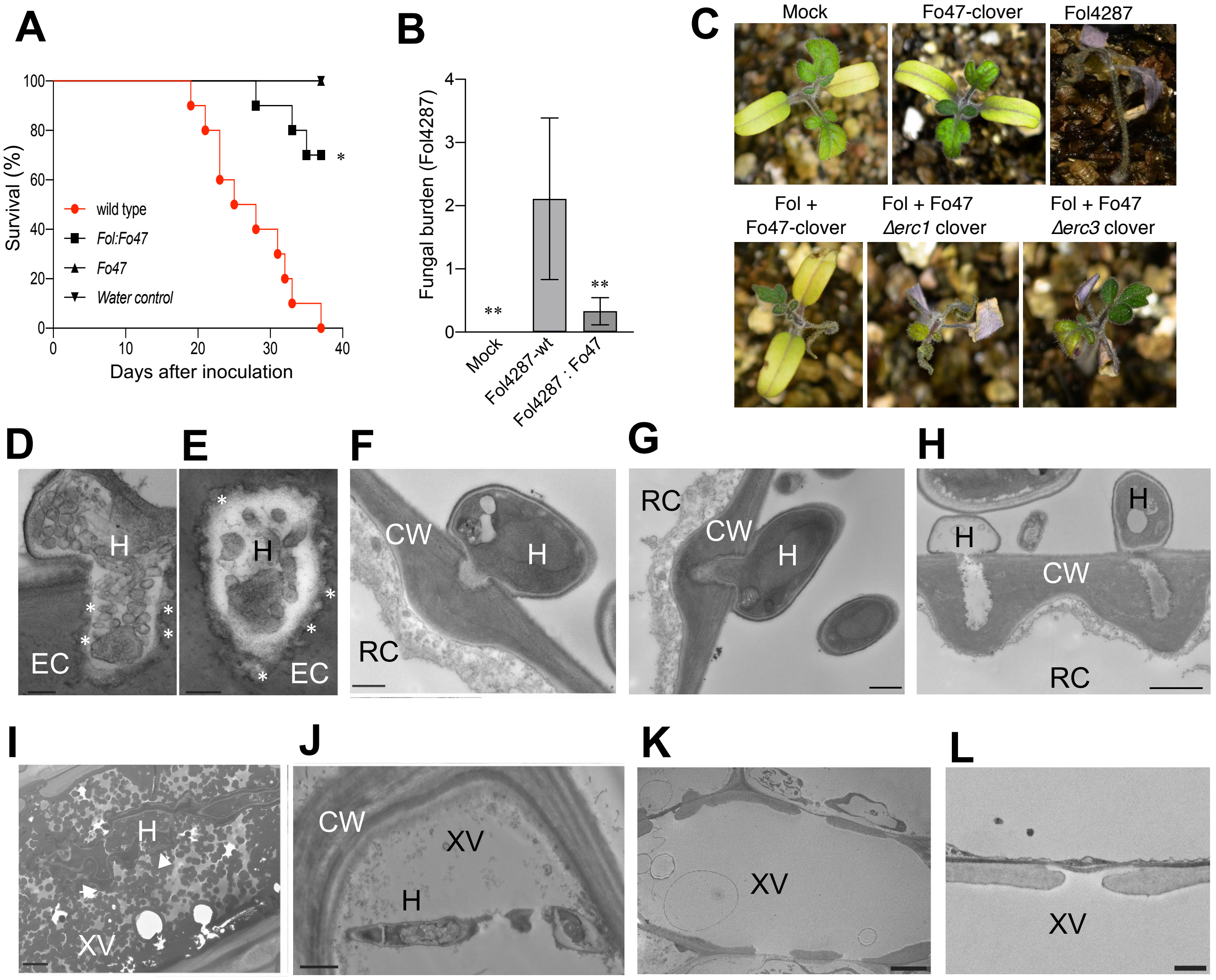

**Fig. S5. ERCs contribute to root colonization and biocontrol ability of Fo47.** (**A**) Kaplan-Meier plot showing the survival of tomato plants inoculated with isolates Fol4287 or Fo47 either individually or together. Number of independent experiments = 3; 10 plants/treatment. Data shown are from one representative experiment. **P* <0.05, versus Fol4287 alone according to log-rank test. Note that Fo47 does not cause mortality when inoculated alone and reduces mortality caused by Fol4287 upon co-inoculation. (**B**) Fungal burden of Fol4287 in roots of tomato plants inoculated either with Fol4287 alone or together with Fo47 was measured by real time (qRT-PCR) of the Fol4287 *six1* gene using total DNA extracted at 10 dpi. Fol4287 DNA was calculated using the threshold cycle (ΔΔCt) method, normalized to the tomato *gadph* gene and expressed relative to Fol4287 alone. Error bars indicate standard deviation (s.d.); n = 3 biological replicates. Asterisks indicate statistical significance versus Fol4287 (one way ANOVA, Bonferroni´s multiple comparison test, p<0.05). The experiment was performed three times with similar results. (**C**) Representative images showing symptoms of tomato plants inoculated with the indicated fungal strains, either alone or in combination, at 25 dpi. (**D** to **J**) TEM micrographs showing the interaction between Fol4287 or Fo47 and tomato root cells at 3 dpi. (**D** and **E**) Hypha (H) of Fol4287 penetrating a root endodermis cell (EC). Note the presence of tubular membrane structures inside and at the edges of the hypha (asterisks). (**F to H**) Fo47 hypha attempting penetration of a root endodermis cell (EC). (**I**) Fo47 hypha inside a xylem vessel (XV). Note encapsulation by granulous material (arrows). (**J**) Fol4287 hypha growing in a xylem vessel. (**K** and **L**). Xylem vessel of an uninoculated control plant. Note absence of granulous material in (**J** to **L**). Scale bars, 0.2 µm in **D** and **E**; 0.5 µm in **F** to **H**; 2 µm in **I** and **J**; 4 µm in **K**; 1 µm in **L**.

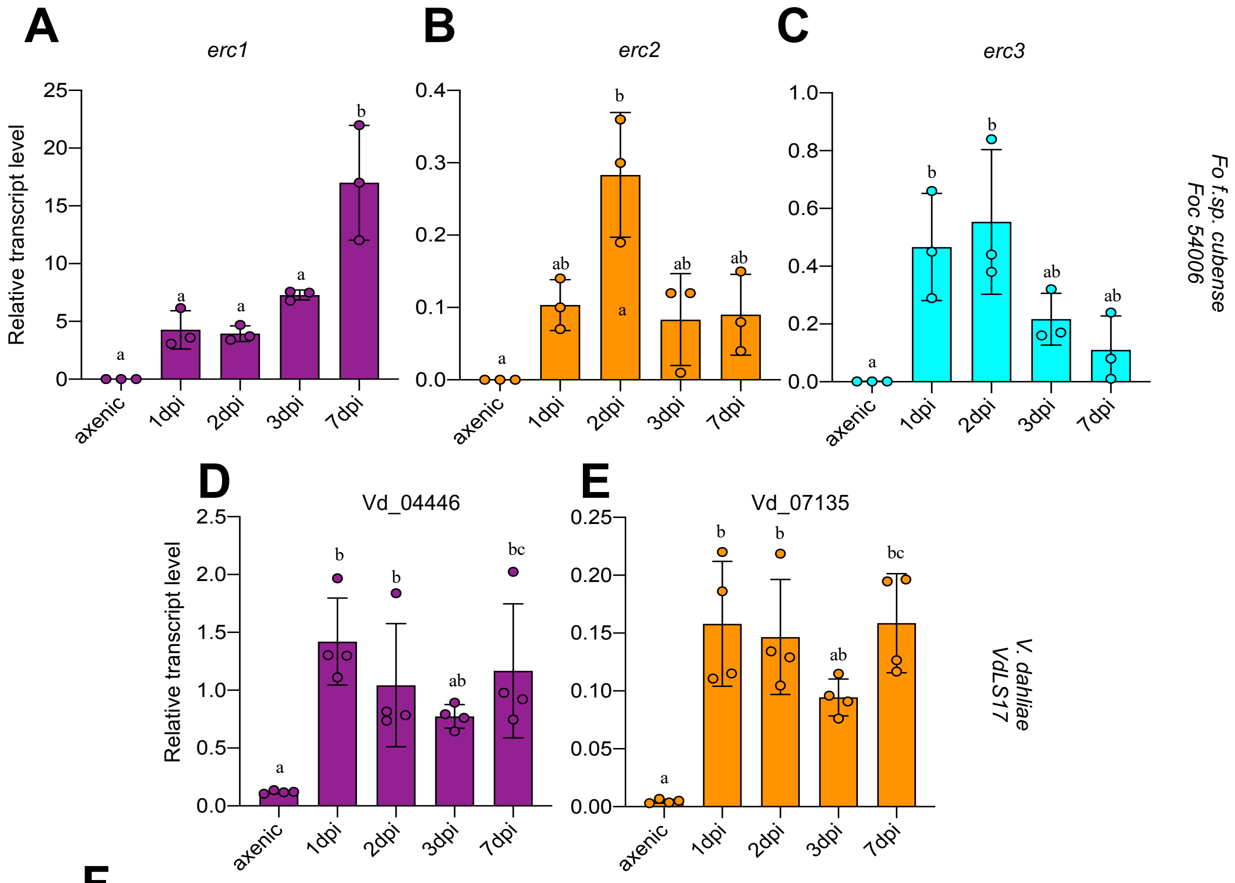

F

| Gene Number | Abbreviation | Homolog Vd | Mol. weight | Signal Peptide | No of cysteines | Predicted domains |
|---|---|---|---|---|---|---|
| FOXG_11583 | ERC1 | VDAG_04446 | 15.8 kDa | Yes | 7 | Lytic transglycolase |
| FOXG_04534 | ERC2 | VDAG_07135 | 32.5 kDa | Yes | 16 | LPMO and Starch binding Domain |

**Fig. S6. ERCs are upregulated during host and non-host interaction in different vascular wilt fungi.** (**A** to **C**) Relative transcript levels of the genes *FOIG_04068* (*erc1*), *FOIG_15314* (*erc2*) and *FOIG_14078* (*erc3)* were measured by qRT-PCR of cDNA obtained from the banana pathogenic isolate Foc54006 grown in minimal medium (axenic) or from roots of tomato plants inoculated with Foc54006 at 1, 2, 3 or 7 dpi. Transcript levels were calculated using the threshold cycle (ΔΔCt) method and normalized to the Fo peptidyl prolyl isomerase (*ppi*) gene. Error bars indicate standard deviation (s.d.); n = 3 biological replicates. Different letters indicate statistically significant differences according to one way ANOVA, Bonferroni´s multiple comparison test ($p < 0.05$). (**D** and **E**) Relative transcript levels of the *erc1 and erc2* homologs were measured by qRT-PCR of cDNA obtained from the isolate VdLS17 of the vascular wilt fungus *Verticillium dahliae* grown in minimal medium (axenic) or from roots of tomato plants inoculated with VdLS17 at 1, 2, 3 or 7 dpi. Transcript levels were calculated using the threshold cycle (ΔΔCt) method and normalized to the *V. dahliae gapdh* gene. Error bars indicate standard deviation (s.d.); n = 3 biological replicates. Different letters indicate statistically significant differences according to one way ANOVA, Bonferroni´s multiple comparison test ($p < 0.05$). (**F**) Table summarizing the characteristics of the ERC1 and ERC2 homologs of *V. dahliae.*

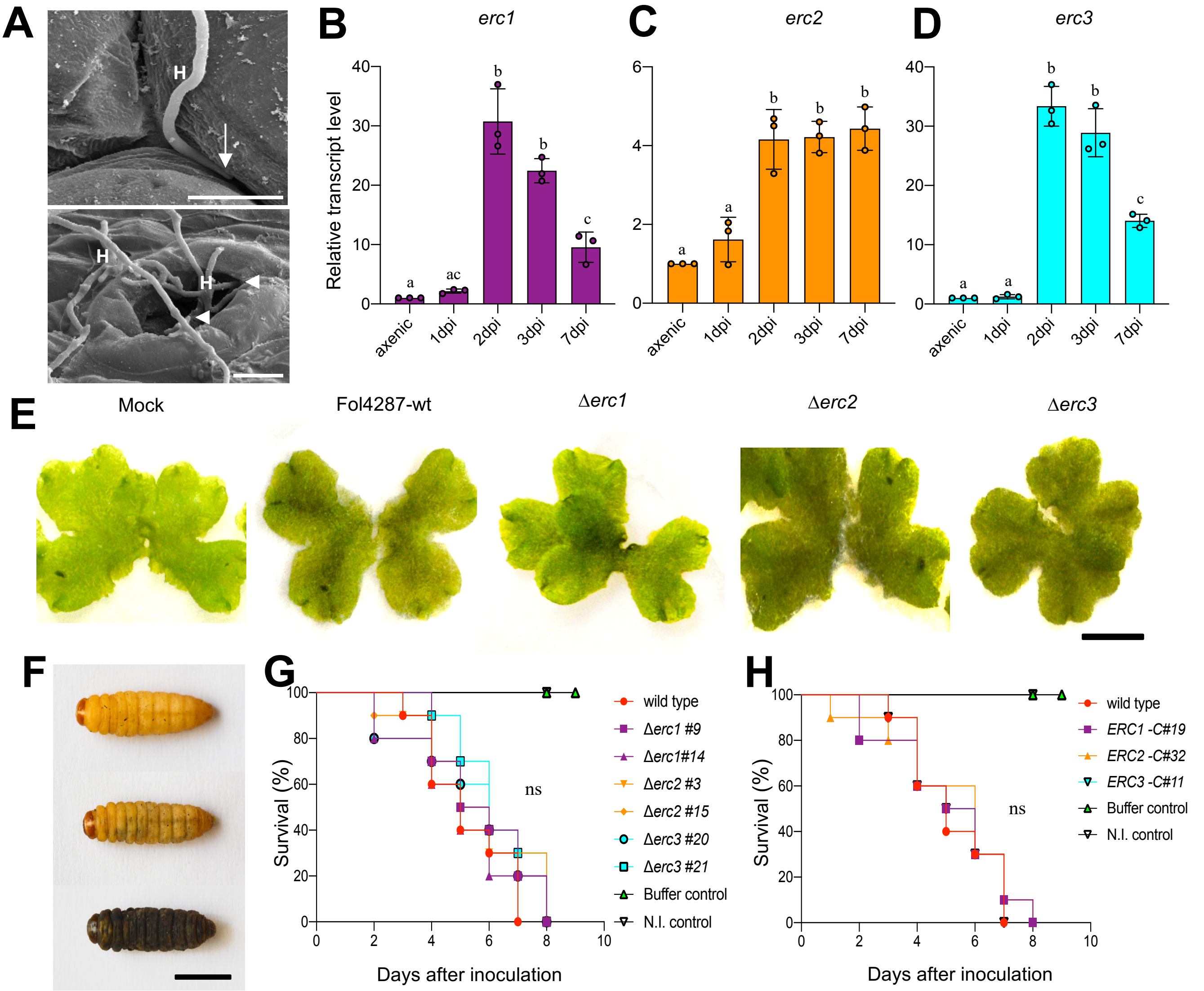

**Fig. S7. ERCs are conserved compatibility factors on vascular and non-vascular plants but not on animal hosts.** (**A**) SEM micrographs showing hyphae (H) entering thalli of *Marchantia polymorpha* intercellularly (arrow) or through open air pores (arrowheads). Scale bars=10 µm. (**B** to **D**) Relative transcript levels of genes *FOXG_11583* (*erc1*), *FOXG_04534* (*erc2*) and *FOXG_16902* (*erc3*) were measured by qRT-PCR of cDNA obtained from Fol4287 grown in minimal medium (axenic) or from Tak-1 thalli inoculated with Fol4287 at 1, 2, 3 or 7 dpi. Transcript levels were calculated using the threshold cycle (ΔΔCt) method and normalized to the Fol4287 peptidyl prolyl isomerase (*ppi*) gene. Error bars indicate standard deviation (s.d.); n = 3 biological replicates. Different letters indicate statistically significant differences according to one way ANOVA, Bonferroni´s multiple comparison test ($p < 0.05$). (**E**) Macroscopic disease symptoms on *M. polymorpha* Tak-1 plants 5 days after dip inoculation with $10^5$ microconidia $ml^{-1}$ of Fol4287, the Δ*erc1*, Δ*erc2 and* Δ*erc3,* deletion mutants or water (mock). Images are representative of three independent experiments. Scale bar, 1 cm. (**F**) Progressive melanisation and killing of *G. mellonella* larvae after inoculation with $1.6 \times 10^4$ microconidia of Fol4287 and incubation at 30°C. (**G** and **H**) Kaplan–Meier plots showing the survival of *G. mellonella* larvae after injection of $1.6 \times 10^4$ microconidia of Fol4287, the Δ*erc1,* Δ*erc2 and* Δ*erc3,* deletion mutants (**G**), their complemented strains (**H**) or buffer control into the hemocoel and incubation at 30°C. Number of independent experiments = 3; 10 larvae/treatment. Data shown are from one representative experiment. ns = non-significant versus Fol4287 according to log-rank test.

### Supplementary Tables

**Supplementary Table 1. Fo reference isolates from the Broad Institute *Fusarium* Comparative Genome Initiative used to test colonization on *Solanum lycopersicum* (Tomato)**

| **Name** | **Description** | **Host plant species** |
|---|---|---|
| NRRL54003 (MN25) | *Fusarium oxysporum* f.sp. *lycopersici* (race 3) | Tomato |
| NRRL54002 (Fo47) | *Fusarium oxysporum* Fo47 (biocontrol strain) | Unknown |
| NRRL54006 (II5) | *Fusarium oxysporum* f.sp. *cubense* (TR4) | Banana |
| NRRL54008 (PHW808) | *Fusarium oxysporum* f.sp. *conglutinans* | Crucifers |
| NRRL54005 | *Fusarium oxysporum* f.sp. *raphani* | Crucifers |
| NRRL26381 (CL57) | *Fusarium oxysporum* f.sp. *radicis- lycopersici* | Tomato |
| NRRL37622 (HDV247) | *Fusarium oxysporum* f.sp. *pisi* | Pea |
| NRRL26406 | *Fusarium oxysporum* f.sp. *melonis* | Melon |
| Fol4287 | *Fusarium oxysporum* f.sp. *lycopersici* (race 2) | Tomato |

**Supplementary Table 2. *Fusarium oxysporum* strains used in the study**

| **Strain** | **Genotype** | **Gene Function** | **Reference** |
|---|---|---|---|
| NRRL34936 (Fol4287) | *Fusarium oxysporum* f.sp. *lycopersici* (race 2) - wildtype | | (ref. 15) |
| NRRL54002 (Fo47) | *Fusarium oxysporum* Fo47 (biocontrol strain) | | (ref. 24) |
| NRRL37622 (HDV247) | *Fusarium oxysporum* f.sp. *pisi* | | (ref. S1) |
| NRRL54003 (MN25) | *Fusarium oxysporum* f.sp. *lycopersici* (race 3) | | (ref. S2) |
| NRRL26381 (CL57) | *Fusarium oxysporum* f.sp. *radicis-lycopersici* | | (ref. S2) |
| NRRL54006 (II5) | *Fusarium oxysporum* f.sp. *cubense (TR4)* | | (ref. S2) |
| NRRL26406 | *Fusarium oxysporum* f.sp. *melonis* | | (ref. S3) |
| NRRL54008 (PHW808) | *Fusarium oxysporum* f.sp. *conglutinans* | | (ref. S2) |
| NRRL54004 | *Fusarium oxysporum* f.sp. *raphani* | | (ref. S2) |
| Fol4287-*mClover3* | *Fol4287::An-gpdaP:*3xFo-*mClover*3:3xFLAG | WT-fluorescent reporter strain | (ref. 47) |
| Fo47-*mClover*3 | *Fo47::An-gpdaP:*3xFo-*mClover*3:3xFLAG | WT-fluorescent reporter strain | (ref. 47) |
| Foc (TR4)-*mClover*3 | *Foc (TR4)::An-gpdaP:*3xFo-*mClover*3:3xFLAG | WT-fluorescent reporter strain | This study |
| *erc1Δ* | *erc1:: HYG* | candidate effector | This study |
| *erc1Δ + erc1* | *erc1:: HYG; erc1:: PHLEO* | candidate effector | This study |
| *erc2Δ* | *erc2:: HYG* | candidate effector | This study |
| *erc2Δ + erc2* | *erc2:: HYG; erc2:: PHLEO* | candidate effector | This study |
| *erc3Δ* | *erc3:: HYG* | candidate effector | This study |
| *erc3Δ + erc3* | *erc3:: HYG; erc3:: PHLEO* | candidate effector | This study |
| *erc4Δ* | *erc4:: HYG* | candidate effector | This study |
| *erc4Δ + erc4* | *erc4:: HYG; erc4:: PHLEO* | candidate effector | This study |
| *erc1Δ erc2Δ* | *erc2:: HYG; erc1:: PHLEO* | candidate effectors | This study |
| *erc2Δ erc3Δ* | *erc2:: HYG; erc3:: PHLEO* | candidate effectors | This study |
| Fo47- *erc1Δ* -*mClover*3 | *Fo47::An-gpdaP:*3xFo-*mClover*3:3xFLAG; *erc1:: PHLEO* | candidate effector | This study |
| Fo47- *erc3Δ* -*mClover*3 | *Fo47::An-gpdaP:*3xFo-*mClover*3:3xFLAG; *erc3:: PHLEO* | candidate effector | This study |
| VdLS17 | *Verticillium dahliae* | | (ref. 46) |

**Supplementary Table 3. Oligonucleotide used in the study.**

| Primer | Sequence | Use | Reference |
|---|---|---|---|
| PHL | AGTTGACCAGTCCGTTCCG | Phleomycin resistance | (ref. 12) |
| LEO | GCCACGAAGTGCACGCAGTT | Phleomycin resistance | (ref. 12) |
| HygY | CGTTGCAAGACCTGCCTGAA | Split marker | (ref. 12) |
| HygG | GGATGCCTCCGCTCGAAGTA | Split marker | (ref. 12) |
| Erc1-P-Fw | CCCAAACGGAATCTCAACAAGCA | erc1 knockout | This study |
| Erc1-P-Rv-Tail | GAATGCACAGGTACACTTGTTTGTAGATGATCGGTGAGTGACCTGTG | erc1 knockout | This study |
| Erc1-T-Fw-Tail | TAGGGGCTGTATTAGGTCTCGGACAGTTGGCAATGGAGGGTT | erc1 knockout | This study |
| Erc1-T-Rv | TCTCGAGCACGGATGGCTTTATA | erc1 knockout | This study |
| Erc1-P-Nested | AACGCGAAGCTGTTCACAAGGA | erc1 knockout | This study |
| Erc1-T-Nested | ATCCCTCTTGAACTTTGAACTCGAG | erc1 knockout | This study |
| Erc2-P-Fw | ATAAGTCCGCCAAGTTAAACCCTG | erc2 knockout | This study |
| Erc2-P-Rv-Tail | GAATGCACAGGTACACTTGTTTGGATCCGTTGATAGCTTTTAGATCTTGA | erc2 knockout | This study |
| Erc2-T-Fw-Tail | TAGGGGCTGTATTAGGTCTCGGGATCCGTTCATTTGAACATCCCAGCATG | erc2 knockout | This study |
| Erc2-T-Rv | TGGCAAGCCGACAAAGGCCAT | erc2 knockout | This study |
| Erc2-P-Nested | ATGGCGTTACAGAGGCCTCA | erc2 knockout | This study |
| Erc2-T-Nested | TGGCAAGCCGACAAAGGCCAT | erc2 knockout | This study |
| Erc3-P-Fw | CGCGTCAGCAAATCTTACATCATCCA | erc3 knockout | This study |
| Erc3-P-Rv-Tail | GAATGCACAGGTACACTTGTTTAGAGCCTTTGACAGAAGATTGAAGAAG | erc3 knockout | This study |
| Erc3-T-Fw-Tail | TAGGGGCTGTATTAGGTCTCGGAGATAGTCTGTTTGAGGAAGCTGGT | erc3 knockout | This study |
| Erc3-T-Rv | TCGTGGAGGATTACCGACAATG | erc3 knockout | This study |
| Erc3-P-Nested | CCTGTCATCGCAAACATTGATCCC | erc3 knockout | This study |
| Erc3-T-Nested | TTGGACTCTTCGCGACGTTC | erc3 knockout | This study |

| | | | |
|---|---|---|---|
| Erc4-P-Fw | TCGATATCACCAACAGAGCCG | erc4 knockout | This study |
| Erc4-P-Rv-Tail | GAATGCACAGGTACACTTGTTTCTTGATAGATAGTTATGTTTGAGTTG | erc4 knockout | This study |
| Erc4-T-Fw-Tail | GTAGGGGCTGTATTAGGTCTCGGGCGATACGTCCGTGACGT | erc4 knockout | This study |
| Erc4-T-Rv | GAAAACAGTCGATCAAGGTATCGACT | erc4 knockout | This study |
| Erc4-P-Nested | CAAGGACTACGTTGCACCGG | erc4 knockout | This study |
| Erc4-T-Nested | TCCAAGGACAGTGGTTTCCG | erc4 knockout | This study |
| gpdA15B | AATAGTGGTGAAATTGATCGTGT | HyG casette/Fo-*mClover* | (ref. 60) |
| Trpc8B | GGATCCAAACAAGTGTACCTGTGCATTC | HyG casette | (ref. 60) |
| SV40 Rev | CTTTTGCTGGCCTTTTGCTCA | Fo-*mClover* | This study |
| act-q7 | ATGTCACCACCTTCAACTCCA | Real time qPCR primer (*actin1*) | (ref. 12) |
| act-q8 | CTCTCGTCGTACTCCTGCTT | Real time qPCR primer (*actin1*) | (ref. 12) |
| gadph-1 | TGATTTGAACTCGTCGCAG | Real time qPCR primer (*SlGAPDH*) | (ref. 12) |
| gadph-2 | CCAAAAACAGTAACAGTAACAGCCTTC | Real time qPCR primer (*SlGAPDH*) | (ref. 12) |
| six-1-1 | ATAGCATGGTACTCCTTGGCG | Real time qPCR primer (*Six1*) | (ref. 12) |
| six-1-2 | CCTGATGGTGACGGTTACGAA | Real time qPCR primer (*Six1*) | (ref. 12) |
| Fo-ppi Fw | AAGGGTGACCAGTTCGATAG | Real time qPCR Primer (*ppi*) | (ref. 47) |
| Fo-ppi Rv | TTCTCGCCGAGCTTCATTTG | Real time qPCR Primer (*ppi*) | (ref. 47) |
| Fo47_10856_Fw | GTTTGGTTGGTTTGGCTCCC | Real time qPCR primer | This study |
| Fo47_10856_Fw | GCAAGTCGTTCCTGCCCTTT | Real time qPCR primer | This study |
| MpEF1a-Fw | CCGAGATCCTGACCAAGG | Real time qPCR primer | (ref. 64) |
| MpEF1a-Fw | GAGGTGGGTACTCAGCGAAG | Real time qPCR primer | (ref. 64) |
| Sl-PR1-Fw | TCTTGTGAGGCCCAAAATTC | Real time qPCR primer (*PR1*) | (ref. 65) |
| Sl-PR1-Rv | TAGTCTGGCCTCTCGGACA | Real time qPCR Primer (*PR1*) | (ref. 65) |
| Sl-GluA-Fw | GGTCTCAACCGCGACATATT | Real time qPCR Primer (*GluA*) | (ref. 65) |
| Sl-GluA-Rv | CACAAGGGCATCGAAAAGAT | Real time qPCR primer (*GluA*) | (ref. 65) |
| Sl-Chi3-Fw | TGCAGGAACATTCACTGGAG | Real time qPCR primer (*Chi3*) | (ref. 65) |

| | | | |
|---|---|---|---|
| Sl-Chi3-Rv | TAACGTTGTGGCATGATGGT | Real time qPCR primer (*Chi3*) | (ref. 65) |
| Sl-Actin-Fw | GAAATAGCATAAGATGGCAG ACG | Real time qPCR primer (*Actin1*) | (ref. 65) |
| Sl-Actin-Rv | ATACCCACCATCACACCAGTA T | Real time qPCR primer (*Actin1*) | (ref. 65) |
| Fo-Erc1-qF | AGCTGGAGACTGATTTCGTG | Real time qPCR primer (*erc1*) | This study |
| Fo-Erc1-qR | TCTTGACGGGGAGGTTACTG | Real time qPCR primer (*erc1*) | This study |
| Fo-Erc2-qF | GTTACAGTCCTGACTGTTCTGC | Real time qPCR primer (*erc2*) | This study |
| Fo-Erc2-qR | CATTGTCAACACCACGGCAT | Real time qPCR primer (*erc2*) | This study |
| Fo-Erc2-qR-Fo47 | GCAATAGACGTATAGCACGAG T | Real time qPCR primer (*erc2*) | This study |
| Fo-Erc3-qF | CAGCGACTCAACTGTCAACAC | Real time qPCR primer (*erc3*) | This study |
| Fo-Erc3-qR | GCTTCCATCACCCTTGTTGAC | Real time qPCR primer (*erc3*) | This study |
| VdGAPDH-qF | CGAGTCCACTGGTGTCTTCA | Real time qPCR primer (*gapdh*) | (ref. 66) |
| VdGAPDH-qR | CCCTCAACGATGGTGAACTT | Real time qPCR primer (*gapdh*) | (ref. 66) |
| Vd-Erc1-qF | GTCTGGCACGTCGCCTTTC | Real time qPCR primer (*erc1*) | This study |
| Vd-Erc1-qR | TCGACAGACGGTGTTCCGAA | Real time qPCR primer (*erc1*) | This study |
| Vd-Erc2-qF | ACGTTGCCGTGACCTTCAAC | Real time qPCR primer (*erc2*) | This study |
| Vd-Erc2-qR | CACGGTGTAGGTCCAGAGG | Real time qPCR primer (*erc2*) | This study |
| Vd-Erc3-qF | CCACTTCGTCCTGCTGCAC | Real time qPCR primer (*erc3*) | This study |
| Vd-Erc3-qR | AGGAGAAGCAGCCGCTATTG | Real time qPCR primer (*erc3*) | This study |

**Supplementary Table 4. Protein Sequences of all the ERC homologs from all Fungal Phylum which are included in the phylogenetic tree.**

**ERC1 homologs**

| **ID** | **MycoCosm protein ID** | **Organism (Portal name MycoCosm)** |
|---|---|---|
| Afum1 6595 | 6595 | *Aspergillus fumigatus* Af293 from AspGD |
| Corb 134341 | 134341 | *Colletotrichum orbiculare* 104-T |
| Ffuj 10272 | 10272 | *Fusarium fujikuroi* IMI 58289 |
| Ffuj 12398 | 12398 | *Fusarium fujikuroi* IMI 58289 |
| Fgra 12756 | 12756 | *Fusarium graminearum* v1.0 |
| Fgra 4001 | 4001 | *Fusarium graminearum* v1.0 |
| Foxy Fo47 23231 | 23231 | *Fusarium oxysporum* Fo47 |
| Foxy fsp lyc 9198 | 9198 | *Fusarium oxysporum* f. sp. *lycopersic*i 4287 v2 |
| Foxy fsp cub 3663 | 3663 | *Fusarium oxysporum* f. sp. *cubense* tropical race 4 54006 (II5) |
| Foxy fsp lyc 7721 (ERC1) | 7721 | *Fusarium oxysporum* f. sp. *lycopersici* 4287 v2 |
| Foxy fsp mat 49012 | 49012 | *Fusarium oxysporum* f. sp. *matthiolae* PHW726 |
| Foxy fsp mel 6322 | 6322 | *Fusarium oxysporum* f. sp. *melonis* (FoMelon) NRRL 26406 |
| Foxy fsp pis 291292 | 291292 | *Fusarium oxysporum* f. sp. *pisi* T415 v1.0 |
| Foxy fsp rad 17198 | 17198 | *Fusarium oxysporum* f. sp. *radices-lycopersici* 26381 (CL57) |
| Foxy fsp rad 7487 | 7487 | *Fusarium oxysporum* f. sp. *radices-lycopersici* 26381 (CL57) |
| Foxy fsp vas 4079 | 4079 | *Fusarium oxysporum* f. sp. *vasinfectum* 25433 (Cotton) |
| Fpse 6318 | 6318 | *Fusarium pseudograminearum* CS3096 |
| Fsol 529717 | 529717 | *Fusarium solani* FSSC 5 v1.0 |
| Fsol 57217 | 57217 | *Fusarium solani* FSSC 5 v1.0 |
| Ftri 544471 | 544471 | *Fusarium tricinctum* MPI-SDFR-AT-0044 v1.0 |
| Fver 5918 | 5918 | *Fusarium verticillioides* 7600 v2 |
| Fver 7860 | 7860 | *Fusarium verticillioides* 7600 v2 |
| Mlar 109181 | 109181 | *Melampsora larici-populina* v2.0 |
| Mlin 203422 | 203422 | *Melampsora lini* CH5 |
| Mory 5403 | 5403 | *Magnaporthe oryzae* 70-15 v3.0 |
| Pgra fsp tri 17381 | 17381 | *Puccinia graminis* f. sp. *tritici* Ug99 haplotype A |

| | | |
|---|---|---|
| Pstr 12867 | 12867 | *Puccinia striiformis* f. sp. t*ritici* 104 E137 A- |
| Ptri1 11083 | 11083 | *Puccinia triticina* 1-1 BBBD Race 1 |
| Rsol 4449 | 4449 | *Rhizoctonia solani* AG-1 IB |
| Scom 2642684 | 2642684 | *Schizophyllum commune* H4-8 v3.0 |
| Uvir 815 | 815 | *Ustilaginoidea virens* |
| Vdah 9690 | 9690 | *Verticillium dahliae* VdLs.17 |
| Ztri XP003855245* | XP003855245 | *Zymoseptoria tritici* IPO323 |
| Foxy fsp pis255841 | 255841 | *Fusarium oxysporum* f. sp. *pisi* T415 v1.0 |
| *** NCBI protien ID** | | |

**ERC2 homologs**

| ID | MycoCosm protein ID | Organism (Portal name MycoCosm) |
|---|---|---|
| Aalt 279631 | 279631 | *Alternaria alternata* SRC1lrK2f v1.0 |
| Anid 10445 | 10445 | *fimicolochytrium* jonesii JEL569 v1.0 |
| Aory 7779 | 7779 | *Aspergillus oryzae* RIB40 |
| Aped 645884 | 645884 | *Agrocybe pediades* AH 40210 v1.0 |
| Arab 8079 | 8079 | *Ascochyta rabiei* ArDII |
| Cgra 4592 | 4592 | *Colletotrichum graminicola* M1.001 |
| Chet 1096518 | 1096518 | *Cochliobolus heterostrophus* C5 v2.0 |
| Chig 11762 | 11762 | *Colletotrichum higginsianum* IMI 349063 |
| Clun 19695 | 19695 | *Cochliobolus lunatus* m118 v2.0 |
| Cmel 7491 | 7491 | *Colletotrichum melonis* CBS 134730 |
| Cnav 526012 | 526012 | *Colletotrichum navitas* CBS125086 v1.0 |
| Cpel 133632 | 133632 | *Coprinellus pellucidus* v1.0 |
| Ffuj 10582 | 10582 | *Fusarium fujikuroi* IMI 58289 |
| Fgra 5476 | 5476 | *Fusarium graminearum* v1.0 |
| Foxy Fo47 8596 | 8596 | *Fusarium oxysporum* Fo47 |
| Foxy fsp cub 17792 | 17792 | *Fusarium oxysporum* f. sp. *cubense* tropical race 4 54006 (II5) |
| Foxy fsp lyc 20802 (ERC2) | 20802 | *Fusarium oxysporum* f. sp. *lycopersici* 4287 v2 |
| Foxy fsp lyc 20991 | 20991 | *Fusarium oxysporum* f. sp. *lycopersici* 4287 v2 |
| Foxy fsp mel 9144 | 9144 | *Fusarium oxysporum* f. sp. *melonis* (FoMelon) NRRL 26406 |
| Foxy fsp mel 9348 | 9348 | *Fusarium oxysporum* f. sp. *melonis* (FoMelon) NRRL 26406 |

| | | |
|---|---|---|
| Foxy fsp pis 6774 | 6774 | *Fusarium oxysporum* f. sp. *pisi* HDV247 |
| Fsol 395350 | 395350 | *Fusarium solani* FSSC 5 v1.0 |
| Ftri 513632 | 513632 | *Fusarium tricinctum* MPI-SDFR-AT-0068 v1.0 |
| Fver 20455 | 20455 | *Fusarium verticillioides* 7600 v2 |
| Lmac 2037 | 2037 | *Leptosphaeria maculans* |
| Ncra 3619 | 3619 | *Neurospora crassa* OR74A v2.0 |
| Pcub 88718 | 88718 | *Psilocybe cubensis* v1.0 |
| Pter 6865 | 6865 | *Pyrenophora teres* f. *teres* |
| Ptra 533134 | 533134 | *Phoma tracheiphila* IPT5 v1.0 |
| Ptri 150726 | 150726 | *Pyrenophora tritici-repentis* |
| Vdah 2653 | 2653 | *Verticillium dahliae* VdLs.17 |
| Xdig 516391 | 516391 | *Xylaria digitata* CBS 161.22 v1.0 |

**ERC3 homologs**

| ID | MycoCosm protein ID | Organism (Portal name MycoCosm) |
|---|---|---|
| Foxy fsp lyc 19501(ERC3) | 19501 | *Fusarium oxysporum* f. sp. *lycopersici* 4287 v2 |
| Foxy fsp mel 8381 | 8381 | *Fusarium oxysporum* f. sp. *melonis* (FoMelon) NRRL 26406 |
| Foxy Fo47 3567 | 3567 | *Fusarium oxysporum* Fo47 |
| Foxy fsp con1 2535 | 2535 | *Fusarium oxysporum* f. sp. *conglutinans* race 2 54008 (PHW808) |
| Foxy fsp vas1 14226 | 14226 | *Fusarium oxysporum* f. sp. *vasinfectum* 25433 (Cotton) |
| Foxy fsp pis 409271 | 409271 | *Fusarium oxysporum* f. sp. *pisi* T415 v1.0 |
| Foxy fsp rad 8606 | 8606 | *Fusarium oxysporum* f. sp. *radicis-lycopersici* 26381 (CL57) |
| Foxy fsp rap 7733 | 7733 | *Fusarium oxysporum* f. sp. *raphani* 54005 |
| Foxy fsp cub 14303 | 14303 | *Fusarium oxysporum* f. sp. *cubense* tropical race 4 54006 (II5) |
| Fcom 564504 | 564504 | *Fusarium commune* MPI-SDFR-AT-0072 v1.0 |
| Fred 527619 | 527619 | *Fusarium redolens* MPI-CAGE-AT-0023 v1.0 |
| Fver 11301 | 11301 | *Fusarium verticillioides* 7600 v2 |
| Ffuj 13894 | 13894 | *Fusarium fujikuroi* IMI 58289 |
| Fpse 6057 | 6057 | *Fusarium pseudograminearum* CS3096 |
| Fgra1 4285 | 4285 | *Fusarium graminearum* v1.0 |
| Aapi1 266161 | 266161 | *Aspergillus spinosus* CBS 483.65 v1.0 |
| Tple 110173 | 110173 | *Trichoderma pleuroti* TPhu1 |
| Tvir 48400 | 48400 | *Trichoderma virens* Gv29-8 v2.0 |
| Afum 8513 | 8513 | *Aspergillus fumigatus* Af293 from AspGD |
| Aalt 113597 | 113597 | *Alternaria alternata* ATCC11680 |
| Xcub 321038 | 321038 | *Xylaria cubensis* CBS 116.85 v1.0 |
| D.PMI573 618292 | 618292 | *Diaporthaceae sp.* PMI_573 v1.0 |
| Pmul 479790 | 479790 | *Phoma multirostrata* 7a v1.0 |
| Cful 197599 | 197599 | *Cladosporium fulvum* v1.0 |

| | | |
|---|---|---|
| Clun 128056 | 128056 | *Cochliobolus lunatus* m118 v2.0 |
| Aory 4052 | 4052 | *Aspergillus oryzae* RIB40 |
| Chet 56451 | 56451 | *Cochliobolus heterostrophus* C4 v1.0 |
| Mory 5787 | 5787 | *Magnaporthe oryzae* 70-15 v3.0 |
| Vdah 9580 | 9580 | *Verticillium dahliae* VdLs.17 |
| Abra 4619 | 4619 | *Alternaria brassicicola* |
| Cgra 8292 | 8292 | *Colletotrichum graminicola* M1.001 |
| Fsol 396389 | 396389 | *Fusarium solani* FSSC 5 v1.0 |
| Corb 135472 | 135472 | *Colletotrichum orbiculare* 104-T |
| Sscl 9047 | 9047 | *Sclerotinia sclerotiorum* v1.0 |
| Ptri 152626 | 152626 | *Pyrenophora tritici-repentis* |
| Anid 7231 | 7231 | *Aspergillus nidulans* |
| Tree 55319 | 55319 | *Trichoderma reesei* v2.0 |
| Bcin 6929 | 6929 | *Botrytis cinerea* v1.0 |
| Ncra 3704 | 3704 | *Neurospora crassa* OR74A v2.0 |
| Srei 4311 | 4311 | *Sporisorium reilianum* SRZ2 |
| Mvul 1484035 | 1484035 | *Mycena vulgaris* CBHHK164 v1.0 |
| Gymear 1114431 | 1114431 | *Gymnopus earleae* GB-263.02 v1.0 |
| Pcub 72876 | 72876 | *Psilocybe cubensis* v1.0 |
| Maph 5476 | 5476 | *Moesziomyces aphidis* DSM 70725 |
